# Serotonin regulates mitochondrial biogenesis and function in rodent cortical neurons via the 5-HT_2A_ receptor and SIRT1-PGC-1α axis

**DOI:** 10.1101/497982

**Authors:** Sashaina E. Fanibunda, Sukrita Deb, Babukrishna Maniyadat, Samir Gupta, Noelia Weisstaub, Jay A. Gingrich, Ashok D.B. Vaidya, Ullas Kolthur-Seetharam, Vidita A. Vaidya

## Abstract

Mitochondria in neurons in addition to their primary role in bioenergetics also contribute to specialized functions including regulation of synaptic transmission, Ca^2+^ homeostasis, neuronal excitability and stress adaptation. However, the factors that influence mitochondrial biogenesis and function in neurons remain poorly elucidated. Here, we identify an important role for serotonin (5-HT) as a regulator of mitochondrial biogenesis and function in rodent cortical neurons, via a 5-HT_2A_ receptor-mediated recruitment of the SIRT1-PGC-1α axis, which is relevant to the neuroprotective action of 5-HT. 5-HT increased mitochondrial biogenesis, reflected through enhanced mtDNA levels, mitotracker staining, and expression of mitochondrial genes. This was accompanied by increased cellular ATP levels, basal and maximal respiration, as well as spare respiratory capacity. Mechanistically the effects of 5-HT were mediated via the 5-HT_2A_ receptor and master modulators of mitochondrial biogenesis, SIRT1 and PGC-1α. SIRT1 was required to mediate the effects of 5-HT on mitochondrial biogenesis and function in cortical neurons. *In vivo* studies revealed that 5-HT_2A_ receptor stimulation increased cortical mtDNA and ATP levels, in a SIRT1 dependent manner. In cortical neurons, 5-HT enhanced expression of anti-oxidant enzymes, decreased cellular reactive oxygen species, and exhibited neuroprotection against excitotoxic and oxidative stress, an effect that required SIRT1. These findings identify 5-HT as a novel upstream regulator of mitochondrial biogenesis and function in cortical neurons, and implicate the mitochondrial effects of 5-HT in its neuroprotective action.

## Introduction

Serotonin (5-HT), a phylogenetically ancient molecule, in addition to its multifaceted neurotransmitter function, has been hypothesized to retain “pre-nervous” roles including trophic and morphogen-like actions (1–3). 5-HT exerts potent effects on neuronal plasticity in the developing and mature nervous system, through an influence on neurite outgrowth, synaptogenesis and synaptic plasticity (1, 4–6). 5-HT also evokes trophic factor-like effects on cell proliferation, survival, and differentiation (3, 7).

Mitochondria are highly dynamic organelles that are vital not only for their bioenergetic role and their influence on cell survival, but also sub-serve specialized functions of regulating excitability, synaptic transmission, buffering Ca^2+^ homeostasis, and modulating structural and functional synaptic plasticity in the context of neurons (8–10). Mitochondrial biogenesis and function in neurons are hypothesized to promote cell viability and mediate effective stress adaptation (11). The vital importance of mitochondria in the context of neurons (12, 13), underscores the importance of studying upstream pathways that drive mitochondrial biogenesis and function in neurons.

Mitochondrial biogenesis and function requires the coordinated transcription of nuclear and mitochondrial encoded genes, and is mediated via transcriptional regulators that respond to extracellular and mitochondrial cues (14, 15). Several reports indicate that the NAD^+^-dependent deacetylase SIRT1 and peroxisome proliferator-activated receptor gamma coactivator 1-alpha (PGC-1α) serve as master regulators of mitochondrial biogenesis and function by controlling gene expression (14, 16–19). While SIRT1 and PGC-1α are implicated in neuronal bioenergetics and survival (16), the upstream cues that recruit the SIRT1-PGC-1α axis in neurons remain relatively unexplored. 5-HT has been reported to increase axonal transport of mitochondria in hippocampal neurons (20), and an influence of 5-HT_1F_ receptor activation on mitochondrial biogenesis in dopaminergic neurons has been recently demonstrated (21).

We hypothesized that 5-HT may exert a putative trophic-like action by serving as an upstream regulator of mitochondrial biogenesis in neurons. To test this hypothesis, we used both *in vitro* cortical neuronal cultures and *in vivo* studies, employing pharmacological and genetic perturbation strategies, and identified a hitherto unknown role of 5-HT as a major regulator of mitochondrial biogenesis and function in cortical neurons, via a 5-HT_2A_ receptor-mediated recruitment of the SIRT1-PGC-1α axis. The effects of 5-HT on mitochondrial biogenesis and function play a central role in the ability of 5-HT to promote neuronal survival in response to excitotoxic or oxidative insults. Our results highlight an important function of 5-HT in the modulation of mitochondrial biogenesis and function in neurons, which facilitates stress adaptation and survival.

## Results

### 5-HT regulates mitochondrial biogenesis and function

We examined the influence of the monoamine, 5-HT on mitochondrial mass in cortical neurons *in vitro* by assessing mitotracker staining, which revealed a significant increase in staining intensity in 5-HT treated neurons (Fig. 1A-C). Immunofluorescence intensity measurements for the mitochondrial voltage dependent anion channel (VDAC) demonstrated significantly increased levels following exposure to 5-HT (Fig. 1D, E). Elevated levels of the mitochondrial outer (VDAC) and inner (cytochrome C, Cyt C), membrane marker proteins, also corroborated the increased mitochondrial mass noted following 5-HT treatment (Fig. 1F-H; Fig. S1A-C). Mitochondrial biogenesis is modulated by the expression of nuclear and mitochondrial-encoded genes, which is orchestrated by master regulators such as PGC-1α, nuclear respiratory factor (*Nrf1*) and transcription factor A, mitochondrial (TFAM) that mediate mtDNA replication. PGC-1α (Fig. 1G, Fig. S1D) and TFAM (Fig. 1H, Fig. S1E) protein levels were significantly elevated following 5-HT exposure. 5-HT treatment also evoked a dose-dependent upregulation in *Ppargc1a* (Fig. 1I), *Tfam* (Fig. 1J) and *Nrf1* (Fig. S1F) transcript expression. A dose-dependent increase in mtDNA content (Fig. 1K) confirmed enhanced mitochondrial biogenesis in response to 5-HT treatment. Time-course analysis (S1G-N) showed a significant increase in PGC-1α expression (Fig. S1G-I), *Tfam* (Fig. S1L, M) and *Nrf1* (Fig. S1L, N) commencing as early as 48 h following onset of 5-HT treatment. This preceded the 5-HT evoked increase in mtDNA (Fig. S1G,J), highlighting the induction of a transcriptional program that may mediate 5-HT-dependent mitochondrial biogenesis.

**Figure 1:**
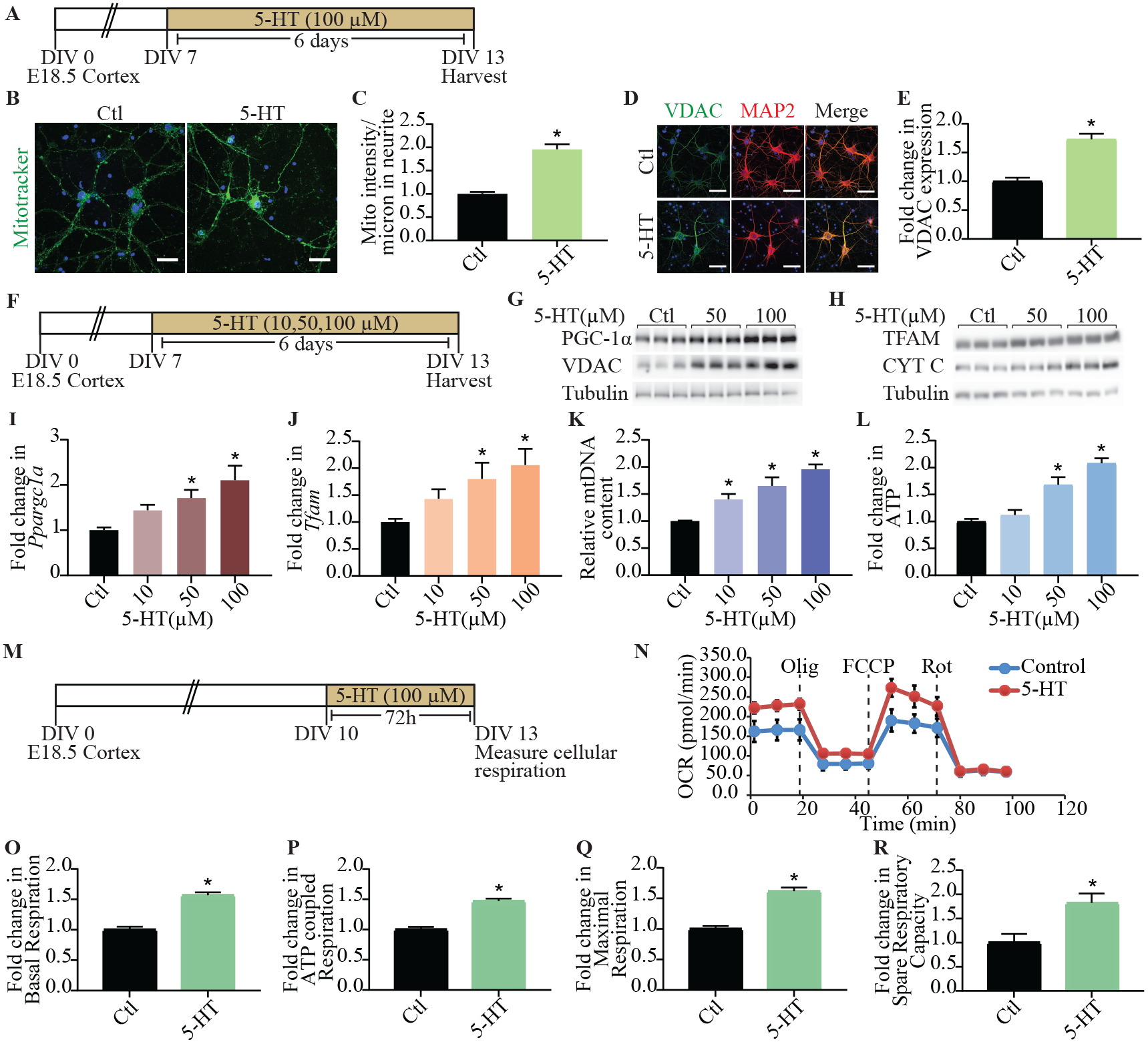
Serotonin regulates mitochondrial biogenesis and function. (A) Shown is a schematic depicting the treatment paradigm with 5-HT (100 μM) in cultured cortical neurons, commencing at day *in vitro* (DIV) 7 until DIV 13. (B) Shown are representative confocal images for mitotracker green staining in control (Ctl) and 5-HT (100 μM) treated neurons, with nuclei counterstained with Hoechst 33342 (blue). Scale bar: 30 μm. Magnification: 60X. (C) Quantitative analysis of fluorescence intensity represented as mitotracker intensity per μm of neurite ± SEM (n = 56 neurons in control, n = 59 neurons for 5-HT treatment, mean values ± SEM compiled across N = 3; \**p* < 0.05 as compared to control, unpaired Student’s *t*-test). (D) Shown are representative immunofluorescence images for VDAC (green), neuronal marker MAP2 protein (red) and merge (yellow) from control and 5-HT (100 μM) treated neurons. Nuclei are counterstained with Hoechst 33342 (blue). Scale bar: 50 μm. Magnification: 60X. (E) Quantitative analysis for VDAC fluorescence intensity are represented as fold change in VDAC expression compared to control ± SEM (n = 86 neurons in control, n = 78 neurons for 5-HT treatment, mean values ± SEM compiled across N = 3; \**p* < 0.05 as compared to control, unpaired Student’s *t-*test). (F) Shown is a schematic depicting treatment of neurons with increasing doses of 5-HT (10, 50, 100 μM). (G and H) Shown are representative immunoblots for PGC-1α, VDAC and tubulin as the loading control (G) and TFAM, Cyt C and tubulin as loading control (H), in neurons treated with increasing doses of 5-HT (50, 100 μM). (I and J) Quantitation of qPCR analysis for mRNA expression of key regulators of mitochondrial biogenesis *Ppargc1a* (I) and *Tfam* (J) are represented as fold change of control ± SEM. (Representative results from n = 4 per treatment group/N = 3, \**p* < 0.05 as compared to control, one-way ANOVA, Tukey’s *post-hoc* test). (K) Quantitative qPCR analysis for mtDNA levels are represented as relative mtDNA content ± SEM. (Representative results from n = 4-6 per treatment group/N = 3, \**p* < 0.05 as compared to control, one-way ANOVA, Tukey’s *post-hoc* test). (L) Quantitation of ATP levels expressed as fold change of control ± SEM. (Representative results from n = 4 per treatment group/N = 4, \**p* < 0.05 as compared to control, one-way ANOVA, Tukey’s *post-hoc* test). (M) Shown is a schematic for the treatment paradigm with 5-HT (100 μM) (DIV 10 - DIV 13) for Seahorse analysis of cellular respiration. (N) Shown is a representative Seahorse plot for 5-HT induced mitochondrial oxygen consumption rate (OCR), with measurements of OCR baseline and following treatment of cells with oligomycin (Olig), FCCP and rotenone (Rot) as indicated. (O-R) Quantitative analysis of the effects of 5-HT on basal respiration (O), ATP coupled respiration (P), maximal respiration (Q), and spare respiratory capacity (R) expressed as fold change of control ± SEM. (Compiled results from n = 5 per treatment group/N = 3, \**p* < 0.05 as compared to control, unpaired Student’s *t*-test).

We next sought to determine if 5-HT altered bioenergetics in cortical neurons, and observed that 5-HT treatment results in enhanced cellular ATP levels in a dose- (Fig. 1L) and time-dependent (Fig. S1G, K) manner. These effects appear to be selective to 5-HT, as neither norepinephrine (NE) nor dopamine (DA) influenced mtDNA content (Fig. S1O, P) or ATP (Fig. S1O, Q) levels in cortical neurons. On assaying for oxidative phosphorylation and electron transport chain (ETC) efficiency, by measuring oxygen consumption rate (OCR) (Fig. 1M, N), we found a robust increase in basal OCR (~ 50%) (Fig. 1O) in 5-HT treated cortical neurons, accompanied by an increase in ATP-coupled respiration (Fig. 1P). Treatment with the mitochondrial uncoupler FCCP, revealed a higher maximal respiration (Fig. 1Q) in 5-HT treated cortical neurons, concomitant with an increase in spare respiratory capacity (Fig. 1R), as compared to controls. These findings indicate that in addition to modulating mitochondrial biogenesis, 5-HT also exerts an important regulatory control on oxidative phosphorylation (OXPHOS) and leads to enhanced mitochondrial function in cortical neurons.

### Effects of 5-HT on mitochondrial biogenesis and function are mediated via the 5-HT_2A_ receptor

We carried out pharmacological and genetic perturbation studies to determine the contribution of specific 5-HT receptors to the effects of 5-HT on mitochondria. Cortical neurons express several 5-HT receptors, amongst which the 5-HT_2A_ and 5-HT_1A_ receptors are expressed at high levels (Fig. S2A, B). The 5-HT_2A_ receptor antagonist MDL100,907, completely inhibited the 5-HT-evoked increase in mtDNA content (Fig. 2A, B), and *Ppargc1a* expression. (Fig. 2C). Further, the 5-HT mediated induction of ATP levels was also prevented by treatment with MDL100,907 (Fig. 2D). A role for 5-HT_2A_ receptors in regulation of mitochondrial biogenesis and energetics was further supported by evidence of a dose-dependent increase in mtDNA (Fig. 2E, F), *Ppargc1a* expression (Fig. 2G) and ATP levels (Fig. 2H) following treatment with the 5-HT_2A_ receptor agonist, DOI. In addition to significant increases in mitochondrial biogenesis and function evoked by DOI, a 5-HT_2A_ receptor agonist with hallucinogenic effects, we also noted that a non-hallucinogenic 5-HT_2A_ receptor agonist, lisuride (Fig. S2C) could also increase mtDNA (Fig. S2D), *Ppargc1a* expression (Fig. S2E) and ATP levels (Fig. S2F). Treatment with the 5-HT_1A_ receptor antagonist, WAY100,635, did not alter the 5-HT mediated increase in mtDNA (Fig. S2G, H) and ATP levels (Fig. S2I). To address whether 5-HT_2A_ receptor stimulation with DOI influences mitochondrial respiration we measured OCR (Fig. 2I, J). Cortical neurons treated with DOI exhibited significant increases in both basal OCR (Fig. 2K) and ATP-coupled respiration (Fig. 2L), accompanied by enhanced maximal respiration (Fig. 2M) and spare respiratory capacity (Fig. 2N). Importantly, the effects of 5-HT on mitochondrial biogenesis and OXPHOS were recapitulated by treatment with the 5-HT_2A_ receptor agonist, DOI.

**Figure 2:**
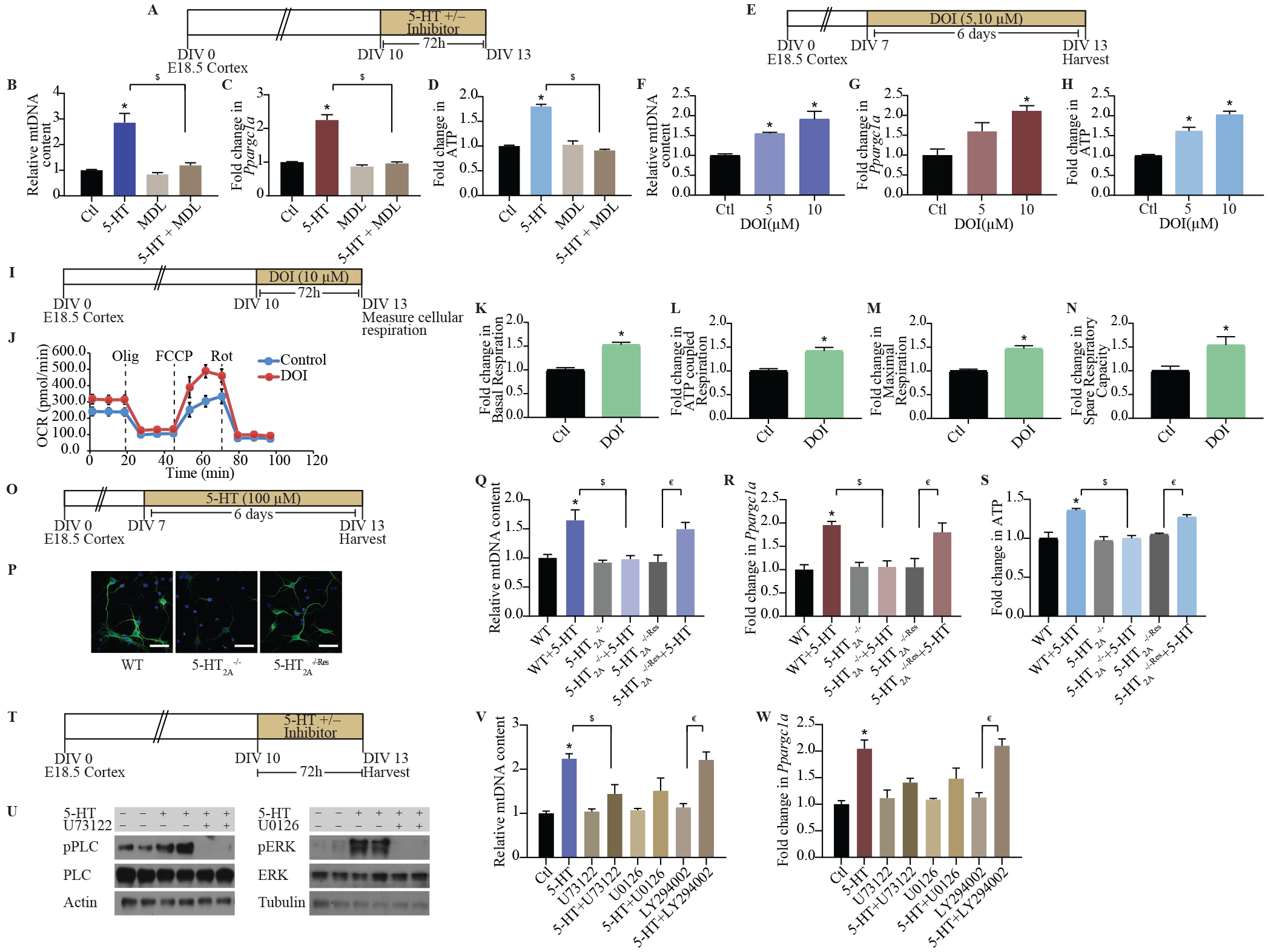
Mitochondrial effects of 5-HT are mediated via the 5-HT_2A_receptor. (A)Shown is a schematic depicting the treatment paradigm for cortical neuron cultures with 5-HT (100 μM), MDL100,907 (10 μM), 5-HT + MDL100,907 for 72 h commencing DIV 10. (B-D) Quantitation of mtDNA (B), *Ppargc1a* mRNA (C) and ATP (D) levels in cortical neurons treated with 5-HT in the presence or absence of MDL100,907 (MDL) are represented as fold change of control (Ctl) ± SEM. (Representative results from n = 4 per treatment group/N = 2 (mtDNA, ATP) N = 3 (*Ppargc1a* mRNA), \**p* < 0.05 as compared to control, ^$^*p* < 0.05 as compared to 5-HT treated group, two-way ANOVA, Tukey’s *post-hoc* test). (E) Shown is a schematic depicting the treatment paradigm for cortical neurons with increasing doses of DOI (5, 10 μM) from DIV 7 to DIV 13. (F-H) Quantitation of mtDNA (F) *Ppargc1a* mRNA (G) and ATP levels (H) in neurons treated with DOI are represented as fold change of control ± SEM. (Representative results from n = 4 per treatment group/N = 2, \**p* < 0.05 as compared to control, one-way ANOVA, Tukey’s *post-hoc* test). (I) Shown is a schematic depicting the treatment paradigm of cortical neurons with DOI (10 μM) for Seahorse analysis of cellular respiration starting DIV 10. (J) Shown is a representative Seahorse plot for control and DOI-treated cortical neurons, with measurements of OCR baseline and following treatment with oligomycin (Olig), FCCP and rotenone (Rot) as indicated. (K-N) Bar graphs depict quantitative analysis of the effects of DOI on basal respiration (K), ATP-coupled respiration (L), maximal respiration (M) and spare respiratory capacity (N) expressed as fold change of control ± SEM. (Compiled results from n = 4-5 per treatment group/N = 3, \**p* < 0.05 as compared to control, unpaired Student’s *t*-test). (O) Shown is a schematic depicting the treatment paradigm of WT, 5-HT_2A_^-/-^ and 5-HT_2A_^-/-Res^ cortical neurons with 5-HT (100 μM). (P) Shown are representative images for 5-HT_2A_ receptor immunofluorescence (green) from WT, 5-HT_2A_^-/-^ and 5-HT_2A_ cortical neuron cultures with nuclei counterstained with Hoechst 33342 (blue). Scale bar: 50 μm. Magnification: 60X. (Q-S) Graphs depict quantitative analysis for mtDNA (Q), *Ppargc1a* mRNA (R) and ATP levels (S) in WT, 5-HT_2A_^-/-^ and 5-HT_2A_^-/-Res^ cortical neurons treated with 5-HT, and results are represented as fold change of WT ± SEM. (Representative results from n = 4 per treatment group/N = 2, \**p* < 0.05 as compared to WT, ^$^*p* < 0.05 as compared to WT+5-HT, ^€^*p* < 0.05 as compared to 5-HT_2A_^-/-Res^, two-way ANOVA, Tukey’s *post-hoc* test). (T)Shown is a schematic depicting the treatment paradigm of cortical neurons with 5-HT (100 μM) in the presence or absence of signaling inhibitors for PLC (U73122, 5 μM), MEK (U0126, 10 μM) and PI3K (LY294002, 10 μM). (U) Shown are representative immunoblots for pPLC and PLC levels in cortical neurons treated with 5-HT in the presence or absence of the PLC inhibitor, U73122, and for pERK and ERK levels in cortical neurons treated with 5-HT in the presence or absence of the MEK inhibitor, U0126. Actin and tubulin immunoblots serve as the loading controls. (V and W) Quantitative analysis for mtDNA (V) and *Ppargc1a* mRNA (W) levels in cortical neurons treated with 5-HT in the presence or absence of PLC, MEK and PI3K inhibitors are represented as fold change of control ± SEM. (Representative results from n = 4 per treatment group/N = 2, \**p* < 0.05 as compared to control, ^$^*p* < 0.05 as compared to 5-HT treated group, ^€^*p* < 0.05 as compared to LY294002 group, two-way ANOVA, Tukey’s *post-hoc* test).

We further characterized the contribution of the 5-HT_2A_ receptor, using cortical neurons derived from 5-HT_2A_ receptor knockouts (5-HT_2A_^-/-^), compared to WT and 5-HT_2A_^-/-Res^ cortical cultures (Fig. 2O, P; Fig. S2J, K). The 5-HT mediated increase in mtDNA content (Fig. 2Q) and *Ppargc1a* expression (Fig. 2R) was completely abrogated in 5-HT_2A_^-/-^ cortical neurons, and was restored in 5-HT_2A_^-/-Res^ cells, wherein a viral-based gene delivery of rAAV8-CaMKIIα-GFP-Cre was utilized to rescue *Htr2a* expression in cortical neurons (Fig. S2J, K). Further, the induction of ATP levels noted following 5-HT treatment to WT neurons was absent in cortical cultures derived from 5-HT_2A_ receptor knockouts, and was reinstated on rescue of 5-HT_2A_ receptor expression (Fig. 2S). Taken together, these results illustrate that the 5-HT_2A_ receptor is necessary for the effects of 5-HT on mitochondrial biogenesis and energetics.

Having identified the 5-HT_2A_ receptor as a key determinant of the effects of 5-HT on mitochondria, we next sought to delineate the contribution of specific downstream signaling pathways. Cortical neurons were incubated with 5-HT in the presence of U73122, U0126 and LY294002, specific inhibitors for the phospholipase C (PLC), MEK and phosphatidylinositol 3-kinase (PI3K) signaling pathways respectively (Fig. 2T, U; Fig. S2L-P). 5-HT treatment resulted in robust activation of pPLC (Fig. 2U, Fig. S2M) and pERK (Fig. 2U, Fig. S2N), but not of pAkt (Fig. S2O, P), which lies downstream of PI3K. The 5-HT mediated increase in mtDNA content (Fig. 2V) and *Ppargc1a* expression (Fig. 2W) were partially blocked by both PLC and MEK inhibitors, but not by PI3K inhibition (Fig. 2V, W). These findings implicate signaling via the PLC and MAPK/ERK cascades in the effects of 5-HT on mitochondrial biogenesis. Collectively, these results indicate that 5-HT via the 5-HT_2A_ receptor, and the PLC and MAP kinase signaling pathways modulate mitochondrial biogenesis.

### Sirt1 is required for the effects of 5-HT on mitochondrial biogenesis and function

SIRT1, a NAD^+^-dependent deacetylase, is well established to induce mitochondrial biogenesis and functions by its ability to regulate transcription involving PGC-1α (14). In 5-HT treated cortical neurons, we observed an upregulation of *Sirt1* mRNA (Fig. 3A, B) and protein (Fig. 3C, D). The transcriptional regulation of *Sirt1* by 5-HT was blocked by co-administration of the 5-HT_2A_ receptor antagonist, MDL100,907 (Fig. S3A, B) and mimicked by the 5-HT_2A_ receptor agonist, DOI (Fig. S3C, D). We investigated the possible involvement of SIRT1 in the actions of 5-HT on mitochondria, by treating cortical neurons with 5-HT in the presence of EX-527, a selective chemical inhibitor of SIRT1 activity (Fig. 3E). EX-527 treatment abrogated the 5-HT-induced increase in mtDNA content (Fig. 3F), and mitochondrial mass in neurites (Fig. 3G, H). Further, co-administration of EX-527 with 5-HT failed to exhibit the 5-HT evoked upregulation of *Ppargc1a* (Fig. 3I), *Tfam* (Fig. 3J), *Nrf1* (Fig. S3E, F) and cytochrome C, *Cycs* (Fig. S3G). We then examined the consequences of SIRT1 inhibition, and found that the 5-HT mediated increase in ATP levels was abrogated in the presence of EX-527 (Fig. 3K). To further ascertain the contribution of SIRT1 to the regulation of ATP levels by 5-HT, we used cortical neurons derived from SIRT1 conditional knockout (Sirt1cKO) embryos (Fig. 3L). 5-HT treatment failed to induce cellular ATP levels in cortical neurons from Sirt1cKO, as compared to wild-type controls, thus demonstrating an essential role for SIRT1 in mediating the effects of 5-HT (Fig. 3L, M). Taken together, our pharmacological and genetic perturbation studies illustrate the role of SIRT1 in mediating the mitochondrial effects of 5-HT.

**Figure 3:**
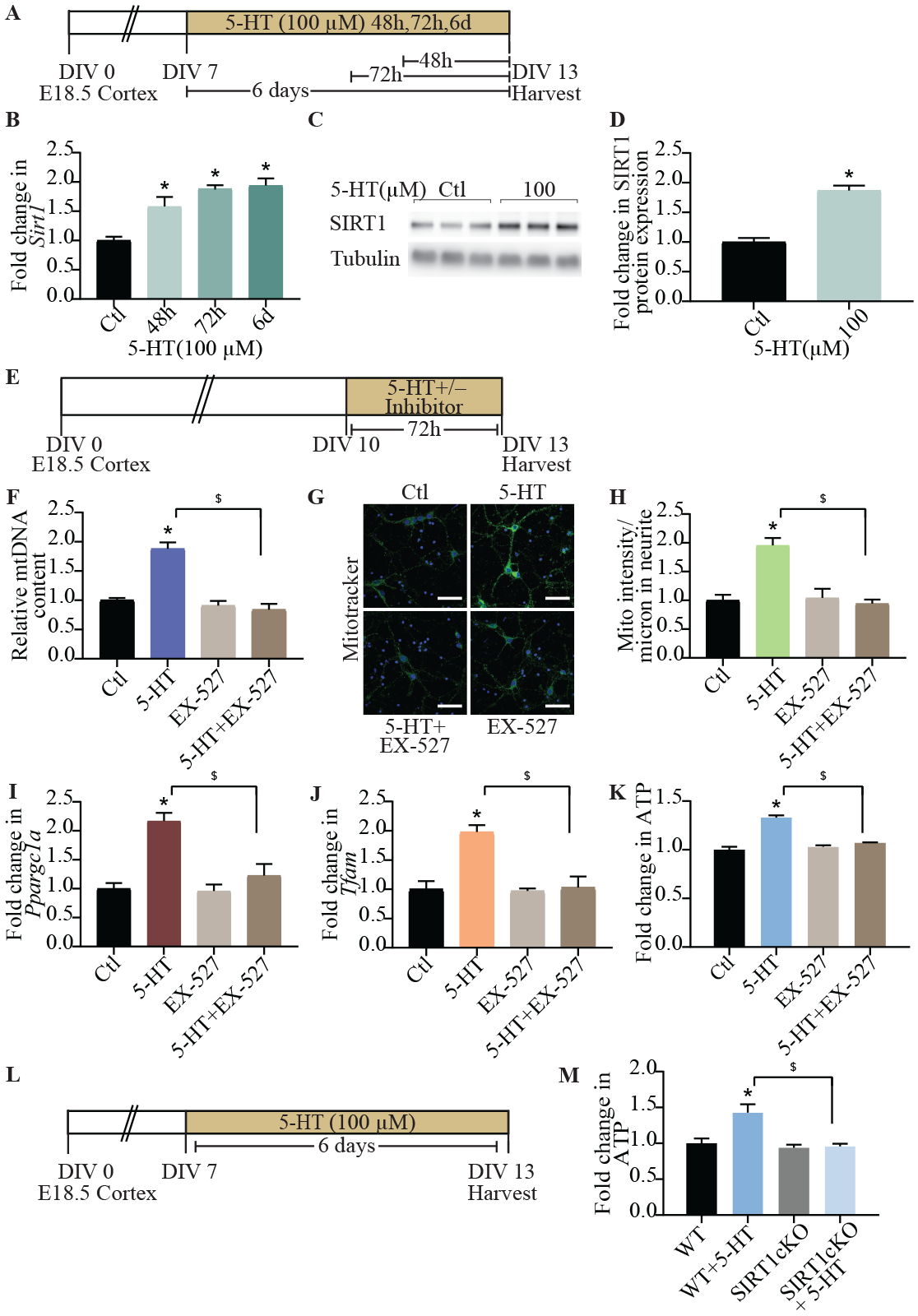
SIRT1 is required for the effects of 5-HT on mitochondria. (A)Shown is a schematic depicting the treatment paradigm of neurons with 5-HT (100 μM) for 48 h, 72 h and 6 days and lysed at DIV 13. (B) Quantitative qPCR analysis for *Sirt1* mRNA expression is expressed as fold change of control (Ctl) ± SEM. (Representative results from n = 4 per treatment group/N = 2, \**p* < 0.05 as compared to control, one-way ANOVA, Tukey’s *post-hoc* test). (C) Shown is a representative immunoblot for SIRT1 protein expression and the loading control tubulin from control and 5-HT (100 μM) treated cortical neurons at 48 h. (D) Quantitative densitometric analysis of SIRT1 levels normalized to tubulin. Results are expressed as fold change of control ± SEM. (Compiled results from n = 6 per treatment group/N = 2, \**p* < 0.05 as compared to control, unpaired Student’s *t*-test). (E) Shown is a schematic depicting the treatment paradigm of neurons with 5-HT (100 μM) in the presence or absence of SIRT1 inhibitor, EX-527 (10 μM) commencing DIV 10. (F) Quantitation of mtDNA in 5-HT treated cortical neurons in the presence or absence of EX-527 are represented as fold change of control ± SEM. (Representative results from n = 4 per treatment group/N = 3, \**p* < 0.05 as compared to control, ^$^*p* < 0.05 as compared to 5-HT treated group, two-way ANOVA, Tukey’s *post-hoc* test). (G) Shown are representative confocal images for mitotracker green staining in control (Ctl) and 5-HT treated cortical neurons in the presence and absence of EX-527. Scale bar: 50 μm. Magnification: 60X. (H) Quantitative analysis of fluorescence intensity represented as mitotracker intensity per μm of neurite ± SEM. (n = 39 neurons in control group, n = 47 neurons in 5-HT treated group, n = 44 neurons in 5-HT+EX-527 group, n = 33 neurons in EX-527 group, mean ± SEM compiled across N = 2; \**p* < 0.05 as compared to control, ^$^p < 0.05 as compared to 5-HT treated group, two-way ANOVA, Tukey’s *post-hoc* test). (I - K) Graphs depict quantitation of *Ppargc1a* (I) and *Tfam* (J) mRNA and ATP (K) levels in cortical neurons treated with 5-HT in the presence or absence of EX-527 and represented as fold change of control ± SEM. (Representative results from n = 4 per treatment group/N = 4 (*Ppargc1a* mRNA and ATP) N = 2 (*Tfam* mRNA), \**p* < 0.05 as compared to control, ^$^*p* < 0.05 as compared to 5-HT treated group, two-way ANOVA, Tukey’s *post-hoc* test). (L) Shown is a schematic depicting the treatment paradigm for cortical neuron cultures from Sirt1cKO and WT mice, with 5-HT (100 μM) commencing DIV 7. (M) Quantitative analysis of ATP levels in WT and SIRTcKO cortical neuron cultures following 5-HT treatment are represented as fold change of WT ± SEM. (Representative results from n = 4 per treatment group/N = 2, \**p* < 0.05 as compared to WT, ^$^*p* < 0.05 as compared to WT+5-HT, two-way ANOVA, Tukey’s *post-hoc* test).

### 5-HT promotes neuroprotective effects against excitotoxic and oxidative stress via the 5-HT_2A_ receptor and SIRT1

Given that 5-HT exerts robust effects on mitochondria, which are major sites of reactive oxygen species (ROS) production and scavenging, we next examined 5-HT effects on cellular ROS levels. We assessed the influence of 5-HT both at baseline and in the context of challenge with excitotoxic (kainate) (Fig. 4A, B) and oxidative (hydrogen peroxide - H_2_O_2_) stressors (Fig. 4A, B). Fluorimetric analysis of cellular ROS levels indicated a baseline reduction following 5-HT treatment (Fig. 4C, D) as compared to control cortical neurons. Further, 5-HT pretreatment robustly attenuated the increased ROS levels observed following treatment with kainate (Fig. 4B, C) or H_2_O_2_ (Fig. 4B, D). Upregulation of ROS scavenging enzymes, superoxide dismutase 2 (*Sod2*) and catalase (*Cat*), which are known to be regulated by SIRT1/PGC-1α (22, 23), corroborated these findings and suggests putative mechanisms for 5-HT-mediated reduction of cellular ROS (Fig. S4A-C).

**Figure 4:**
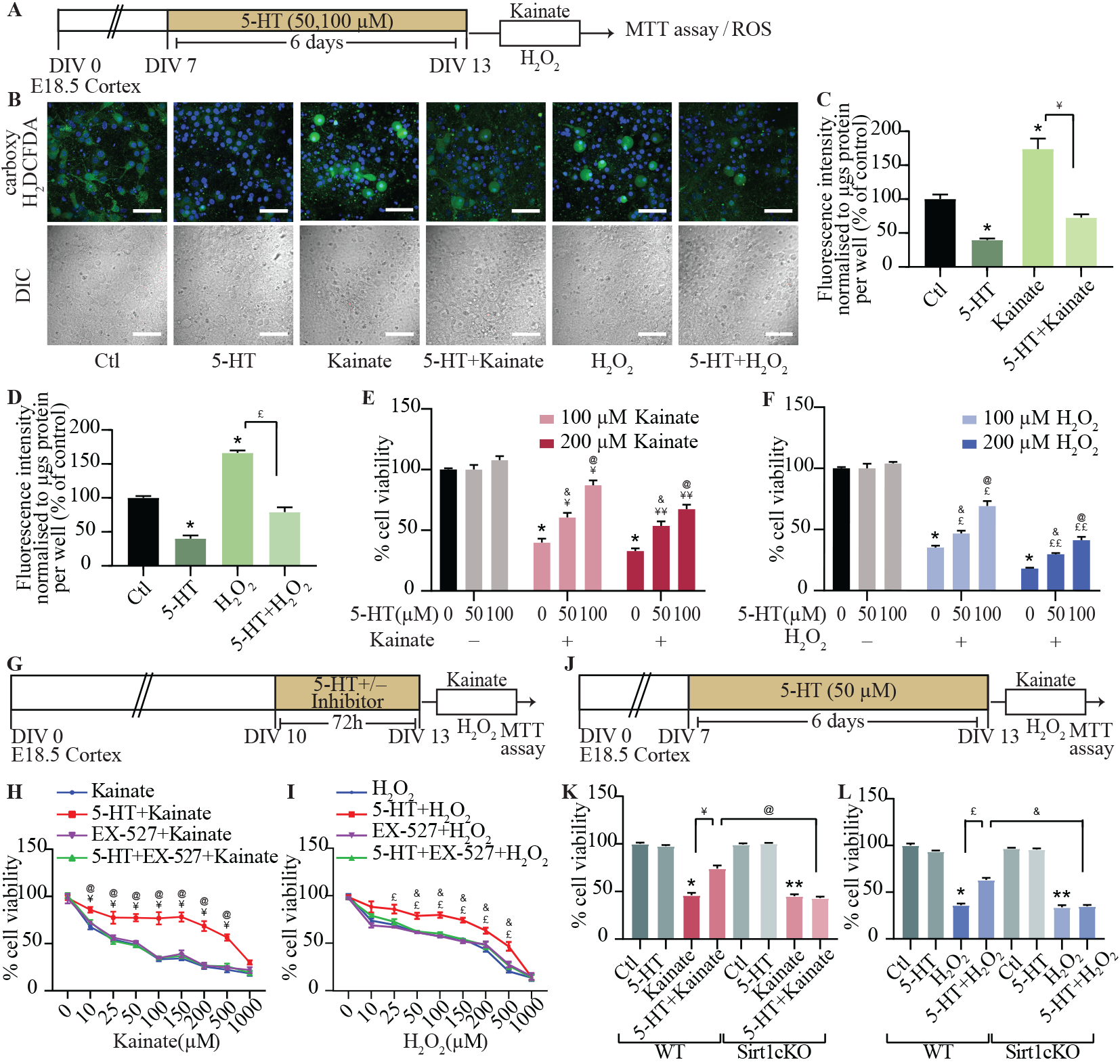
5-HT exerts neuroprotective effects against excitotoxic and oxidative stress via SIRT1. (A) Shown is a schematic depicting the treatment paradigm of neurons with 5-HT (50 or 100 μM) pretreatment commencing DIV 7 followed by exposure to the excitotoxic insult of kainate (100, 200 μM) or oxidative stress through H_2_O_2_ (100, 200 μM) on DIV 13, and analysis of cellular reactive oxygen species (ROS) or cell survival via the MTT assay. (B) Shown are representative confocal images of carboxy-H_2_DCFDA staining (green) to measure cellular ROS in neurons treated with 5-HT (100 μM), kainate (100 μM), H_2_O_2_ (200 μM), 5-HT + kainate, 5-HT + H_2_O_2_. Nuclei are counterstained with Hoechst 33342 (blue). Shown are the DIC images of cortical neurons across treatment conditions. Scale bar: 50 μm. Magnification: 60X. (C and D) Fluorimetric quantitation of staining intensity of cellular ROS (carboxy-H_2_DCFDA) normalized to protein content per well and represented as percent of control ± SEM. (Representative results from n = 4 per treatment group/N = 2, \**p* < 0.05 as compared to control, ^¥^*p* < 0.05 as compared to kainate treated group, ^£^*p* < 0.05 as compared to H_2_O_2_ treated group, two-way ANOVA, Tukey’s *post-hoc* test). (E and F) Graphs depict cell viability assessed by the MTT assay in cortical neurons challenged with kainate (100 μM or 200 μM) or H_2_O_2_ (100 μM or 200 μM) with or without pretreatment with 5-HT (50 μM or 100 μM). Results are expressed as percent of control cell viability ± SEM. (Representative results from n = 3 per treatment group/N = 2, \**p* < 0.05 as compared to control, ^¥^*p* < 0.05 as compared to 100 μM kainate treated group, ^¥¥^*p* < 0.05 as compared to 200 μM kainate treated group, ^£^*p* < 0.05 as compared to 100 μM H_2_O_2_ treated group, ^££^*p* < 0.05 as compared to 200 μM H_2_O_2_ treated group, ^&^*p* < 0.05 as compared to 50 μM 5-HT treated group, ^@^*p* < 0.05 as compared to 100 μM 5-HT treated group, two-way ANOVA, Tukey’s *post-hoc* test). (G) Shown is a schematic depicting the treatment paradigm of neurons with 5-HT (100 μM) in the presence or absence of the SIRT1 inhibitor, EX-527 (10 μM), followed by challenge with increasing doses of kainate or H_2_O_2_ (0 μM to 1000 μM) and analysis of cell viability using the MTT assay. (H) Shown is a line graph for cell viability of cortical neurons in response to increasing doses of kainate (0 μM to 1000 μM) with treatment groups of kainate (blue), 5-HT + kainate (red), EX-527 + kainate (purple), 5-HT + EX-527 + kainate (green), expressed as per cent of control cell viability ± SEM. (Representative results from n = 3 per treatment group/N = 2, ^¥^*p* < 0.05 as compared to kainate treated group, ^@^*p* < 0.05 as compared to 5-HT + EX-527 + kainate treated group, three-way ANOVA, Tukey’s *post-hoc* test). (I) Shown is a line graph for cell viability of cortical neurons in response to increasing doses of H_2_O_2_ (0 μM to 1000 μM) with treatment groups of H_2_O_2_ (blue), 5-HT + H_2_O_2_ (red), EX-527 + H_2_O_2_ (purple), 5-HT + EX-527 + H_2_O_2_ (green), expressed as per cent of control cell viability ± SEM. (Representative results from n = 3 per treatment group/N = 2, ^£^*p* < 0.05 as compared to H_2_O_2_ treated group, ^&^*p* < 0.05 as compared to 5-HT + EX-527 + H_2_O_2_ treated group, three-way ANOVA, Tukey’s *post-hoc* test). (J) Shown is a schematic depicting the treatment paradigm of cortical neuron cultures derived from WT and Sirt1cKO embryos, with 5-HT (50 μM) treatment commencing DIV 7 followed by a challenge with kainate (100 μM) or H_2_O_2_ (200 μM) on DIV 13 and analysis of cell viability. (K and L) Graph depicts cell viability of WT, WT + 5-HT, Sirt1cKO, Sirt1cKO + 5-HT cortical neurons left untreated (Ctl) or challenged with kainate (K) or H_2_O_2_ (L), and expressed as per cent of WT-Ctl cell viability ± SEM. (Representative results from n = 3 per treatment group/N = 2, \**p* < 0.05 as compared to WT-Ctl, \*\**p* < 0.05 as compared to Sirt1cKO-Ctl, ^¥^*p* < 0.05 as compared to WT+kainate, ^@^*p* < 0.05 as compared to WT+5-HT+kainate, ^£^*p* < 0.05 as compared to WT+H_2_O_2_, ^&^*p* < 0.05 as compared to WT+5-HT+H_2_O_2_, three-way ANOVA, Tukey’s *post-hoc* test).

We then sought to examine potential neuroprotective effects of 5-HT in the context of kainate-mediated excitotoxicity or H_2_O_2_-mediated oxidative damage (Fig. 4A). Cell survival assays revealed that cortical neuronal survival was significantly reduced by kainate (Fig. 4E) and H_2_O_2_ (Fig. 4F) treatment, and was attenuated in a dose-dependent fashion in 5-HT pretreated cortical cultures. These neuroprotective effects of 5-HT were mimicked by pretreatment with DOI, which also enhanced cell survival to excitotoxic or oxidative insults (Fig. S4D-F). Given our results of a role for SIRT1 in the mitochondrial effects of 5-HT, we examined the contribution of SIRT1 to the neuroprotective action of 5-HT. Cortical neuronal cultures were pretreated with 5-HT in the presence/absence of the SIRT1 inhibitor, EX-527, followed by exposure to increasing doses of either kainate (Fig. 4G, H) or H_2_O_2_ (Fig. 4G, I). EX-527 prevented the 5-HT-mediated neuroprotective effects in response to challenge with escalating doses of kainate (Fig. 4H) and H_2_O_2_ (Fig. 4I). 5-HT exerted neuroprotective actions over a wide-range of kainate and H_2_O_2_ doses, and SIRT1 inhibition prevented these neuroprotective actions of 5-HT. The enhanced cell viability noted in cortical neurons pretreated with the 5-HT_2A_ receptor agonist, DOI, on kainate or H_2_O_2_ challenge, was attenuated by EX-527 (Fig. S4G-I). These findings indicate that SIRT1 plays an important role in contributing to the neuroprotective effects of 5-HT, and the 5-HT_2A_ receptor agonist, DOI, in the context of excitotoxic and oxidative stress. This is further confirmed via evidence from Sirt1cKO cortical cultures, which unlike WT cortical neurons fail to exhibit the improved cell viability noted on 5-HT pretreatment prior to kainate (Fig. 4J, K) or H_2_O_2_ (Fig. 4J, L) mediated challenge. Taken together, these results indicate that 5-HT via the 5-HT_2A_ receptor and the sirtuin, SIRT1, exerts robust neuroprotective effects against excitotoxic and oxidative cell death and damage.

### 5-HT_2A_ receptor mediated in vivo regulation of cortical mitochondrial DNA content, gene expression and ATP requires SIRT1

Given we noted robust effects of 5-HT, via the 5-HT_2A_ receptor, on mitochondrial biogenesis and function in cortical neurons *in vitro*, we next sought to examine whether DOI-mediated stimulation of the 5-HT_2A_ receptor led to regulation of mitochondrial biogenesis, gene expression and cellular ATP levels *in vivo* (Fig. 5A). Sub-chronic (72 h) treatment with DOI resulted in a significant increase in mtDNA content (Fig. 5B) as well as expression of *Ppargc1a*, *Sirt1*, *Nrf1, Tfam*, and *Cycs* levels (Fig. 5C) in the rat neocortex. This was accompanied by enhanced cortical cellular ATP levels in DOI treated animals as compared to controls (Fig. 5D). These *in vivo* observations recapitulated the effects of DOI on mitochondria noted in cortical neuronal cultures.

**Figure 5:**
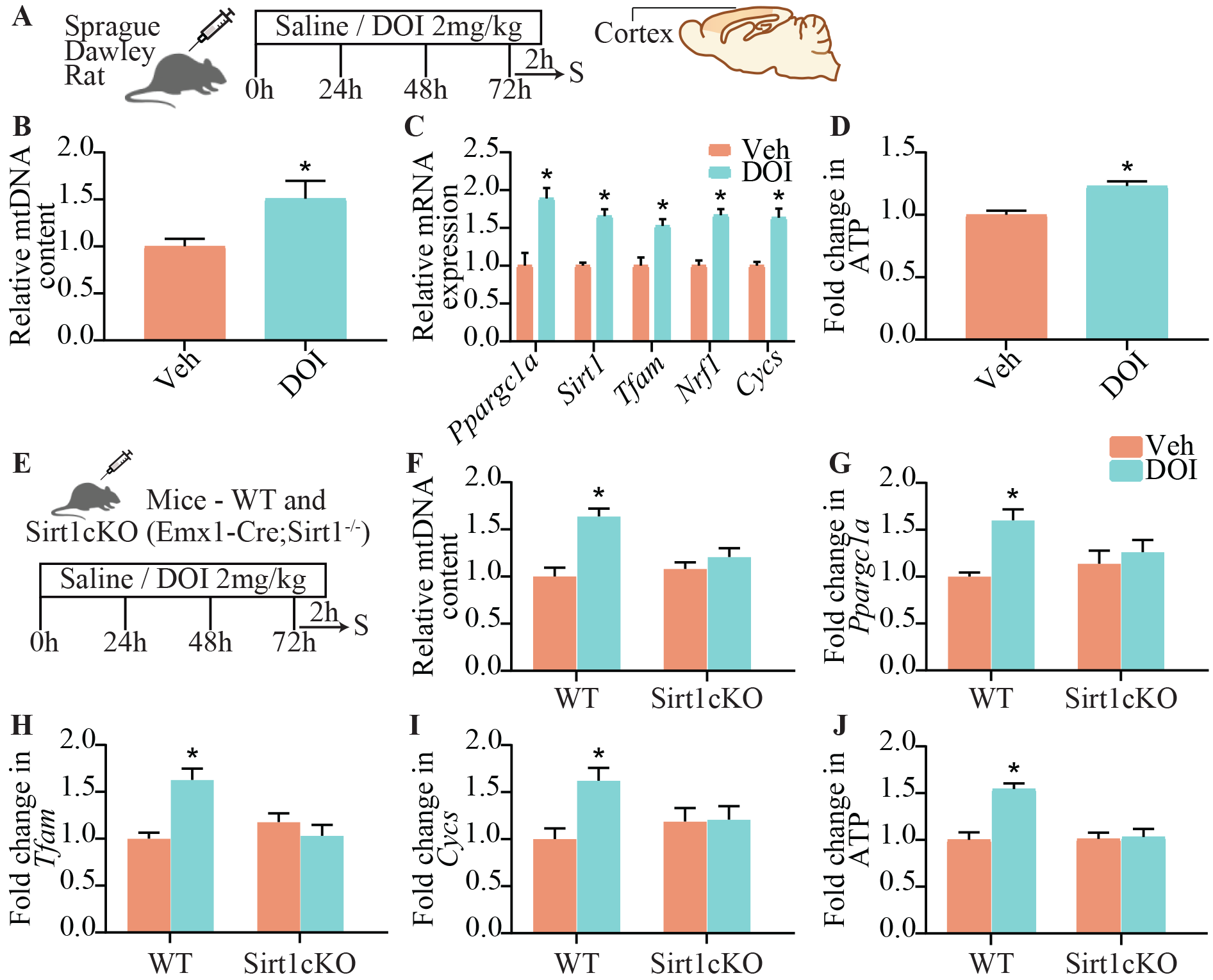
Serotonin_2A_ receptor mediated regulation of cortical mitochondrial DNA content, gene expression and ATP in in vivo requires Sirt1. (A)Shown is a schematic of the treatment paradigm for Sprague-Dawley rats with the 5-HT_2A_ receptor agonist, DOI (2 mg/kg) or vehicle (saline). (B-D) Quantitation of mtDNA levels (B), gene expression of *Ppargc1a*, *Sirt1*, *Tfam*, *Nrf1*, *Cycs* (C) and ATP levels (D) within the cortex of saline and DOI treated rats expressed as fold change of vehicle ± SEM. (n = 8 to 11 animals per group, \**p* < 0.05 as compared to vehicle, unpaired Student’s *t*-test). (E)Shown is a schematic of the treatment paradigm for WT and Sirt1cKO mice with the 5-HT_2A_ receptor agonist, DOI (2 mg/kg) or vehicle (saline). (F-J) Quantitation of mtDNA levels (F), mRNA expression of *Ppargc1a* (G), *Tfam* (H) and *Cycs* (I) and ATP levels (J), in WT and SIRT1cKO mice treated with DOI are expressed as fold change of WT+Veh ± SEM. (n = 8 to 12 animals per group, \**p* < 0.05 as compared to WT, two-way ANOVA, Tukey’s *post-hoc* test).

We then addressed the contribution of SIRT1 to the effects of 5-HT_2A_ receptor stimulation on mtDNA, gene expression of mitochondrial regulators and ATP levels *in vivo* using conditional, cortical SIRT1 knockout (Sirt1cKO: Emx1-Cre; Sirt1^-/-^) mice (Fig. 5E). While sub-chronic DOI administration resulted in a significant induction of mtDNA levels in wild-type (WT) animals, this was lost in Sirt1cKO mice (Fig. 5F). Further, DOI treatment evoked an upregulation of *Ppargc1a*, *Tfam*, and cytochrome C, *Cycs*, in WT, but not in Sirt1cKO mice (Fig. 5G-I). Furthermore, the DOI-mediated increase in cellular ATP levels was completely abrogated in Sirt1cKO mice (Fig. 5J). Collectively, these results establish that 5-HT_2A_ receptor stimulation induces mitochondrial biogenesis and function through transcriptional control exerted by SIRT1.

## Discussion

Our findings demonstrate that 5-HT regulates both mitochondrial biogenesis and function in cortical neurons, via the 5-HT_2A_ receptor. 5-HT_2A_ receptor stimulation recruits the SIRT1-PGC-1α axis, through PLC and MAPK signaling pathways, which enhance transcription of key factors such as NRF1 and TFAM that drive mitochondrial biogenesis. We show that SIRT1 is essential for the mitochondrial effects of 5-HT on cortical neurons. Further, 5-HT enhances ATP production and OXPHOS, accompanied by increased basal and maximal respiration, and improved spare respiratory capacity, an effect also recapitulated through 5-HT_2A_ receptor stimulation. In agreement with our *in vitro* data, 5-HT_2A_ receptor stimulation *in vivo*, enhanced mitochondrial biogenesis, gene expression of several mitochondrial regulators, and ATP levels within the neocortex, an effect that was dependent on SIRT1. Pre-exposure to 5-HT enhances the ability of cortical neurons to buffer excitotoxic and oxidative stress, accompanied by a reduction in cellular ROS. SIRT1 is required for these pro-survival effects of 5-HT and 5-HT_2A_ receptor stimulation, suggesting the intriguing possibility that the mitochondrial effects of 5-HT may underlie the enhanced stress-adaptation noted in cortical neurons. Our findings indicate that 5-HT, through the 5-HT_2A_ receptor, is an important upstream factor that regulates neuronal mitochondrial biogenesis and function via SIRT1 (Fig. 6). Further, we highlight a robust pro-survival effect of 5-HT and DOI on cortical neurons when challenged with excitotoxic and oxidative insults, an effect mediated via SIRT1.

**Figure 6:**
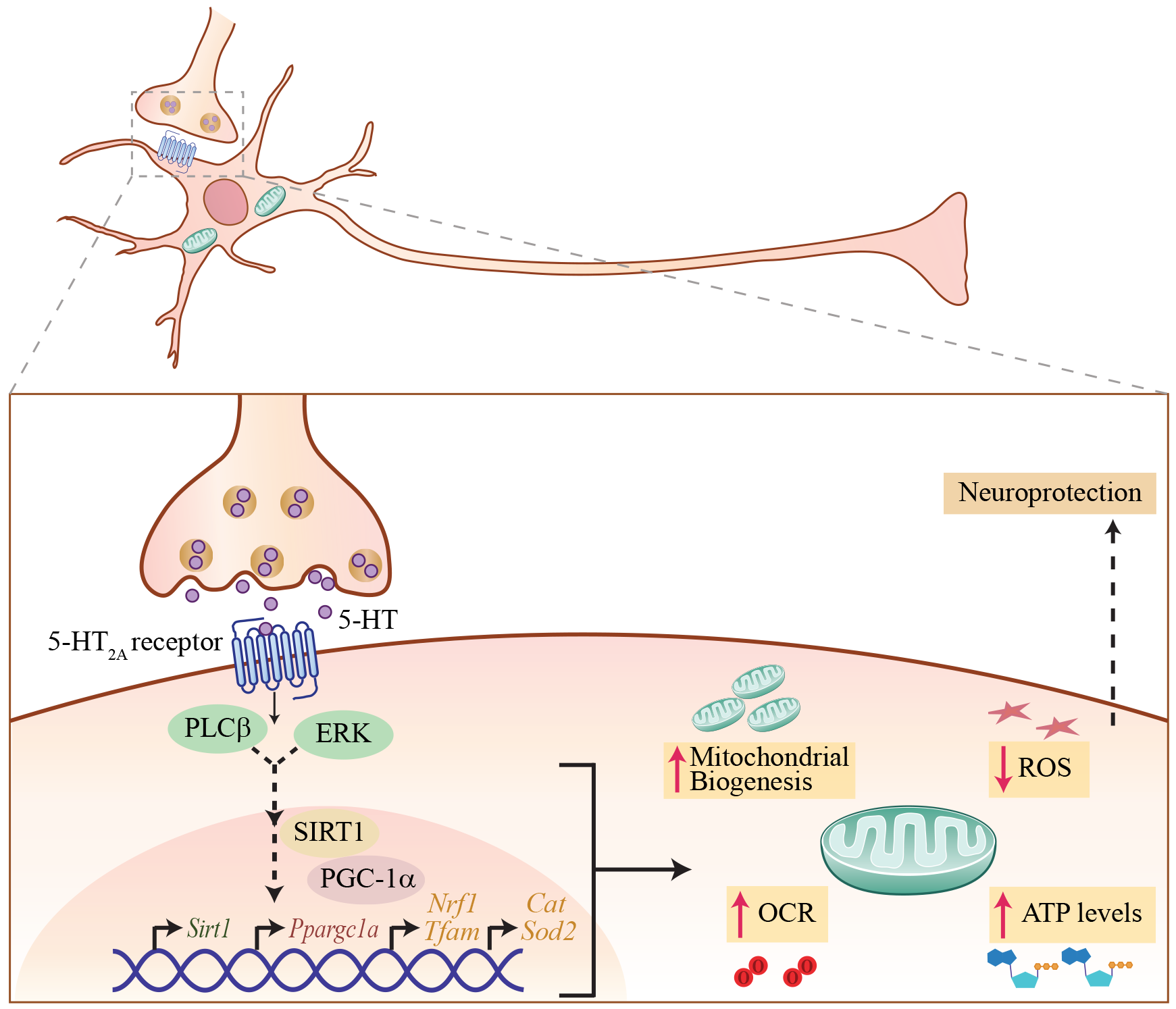
Schematic depicting the putative mechanism for 5-HT effects on mitochondria. (A) Shown is a model depicting a putative mechanism for the effects of 5-HT on mitochondrial biogenesis and function. 5-HT via the 5-HT_2A_ receptor expressed by cortical neurons evokes an activation of the PLC and MAPK signaling pathways. This recruitment of PLC and MAPK signaling, likely through a multiple step process, results in recruitment of SIRT1, which is a key regulator of mitochondrial biogenesis. SIRT1 may then modulate PGC-1α expression which has been shown to be a master regulator of mitochondrial biogenesis and in turn could enhance NRF1 and mitochondrial transcription factor TFAM expression, thus mediating effects of 5-HT on mitochondrial biogenesis. Our results indicate that SIRT1 is required for the effects of both 5-HT and 5-HT_2A_ receptor agonists on mitochondrial biogenesis. Further, 5-HT and 5-HT_2A_ receptor agonists also enhance ATP production and increase OXPHOS, basal and maximal respiration, and spare respiratory capacity. 5-HT also reduces cellular ROS levels and enhances expression of antioxidant enzymes in cortical neurons. This enhancement in mitochondrial biogenesis and function may contribute to the effects of 5-HT, and 5-HT_2A_ receptor agonists, in conferring neuronal protection from excitotoxic and oxidative stress.

Prior reports indicate that specific 5-HT receptors can influence mitochondrial biogenesis in non-neuronal cells, such as renal proximal tubular cells and cardiomyocytes. 5-HT_1F_ and 5-HT_2A_ receptor activation in renal proximal tubular cells can promote mitochondrial biogenesis through recruitment of PGC-1α, and enhance recovery of mitochondrial function following kidney injury (24–27). In cardiomyocytes, 5-HT promotes cell survival via the 5-HT_2B_ receptor, and PI3K-Akt and MAPK pathways (28). Perinatal SSRI exposure is linked to increased mitochondrial respiratory capacity and reduced ROS levels in cardiac tissue (29) and liver (30), as well as sex-specific effects on mitochondrial function within the brainstem (31). Thus far, few studies have examined the direct influence of 5-HT on mitochondria in neurons. 5-HT is reported to promote mitochondrial transport in hippocampal neurons, via the 5-HT_1A_ receptor (20). A recent study demonstrates that 5-HT_1F_ receptor stimulation regulates mitochondrial biogenesis in nigral dopaminergic neurons, and exerts neuroprotective effects in an animal model of Parkinson’s disease (21). Our findings highlight a key role of 5-HT via the 5-HT_2A_ receptor, PLC and MAPK pathways, and the SIRT1-PGC-1α axis in enhancing mitochondrial biogenesis and function both in cortical neurons *in vitro* and in the neocortex *in vivo*. Collectively, these results highlight a putative “pre-nervous” trophic factor-like action of 5-HT on mitochondria, and suggest the possibility that tissue-specific mitochondrial effects of 5-HT may involve distinct 5-HT receptors that facilitate cell survival and stress adaptation.

Interestingly, we did not observe any effect of norepinephrine and dopamine on mitochondrial biogenesis or ATP levels in cortical neurons. Dopamine receptors have been linked to modulation of mitochondrial respiration in the nucleus accumbens (32) and to inhibition of mitochondrial motility in hippocampal neurons (33), suggesting that other monoamines could exert circuit-specific influences on mitochondria. In this vein, it is of interest that melatonin and N-acetylserotonin, a precursor for melatonin, both exhibit neuroprotective effects, hypothesized to involve inhibition of mitochondrial death pathways (34–36). While these reports suggest that diverse monoamines could impinge on mitochondrial mechanisms as a component of their neuroprotective action, they do not address whether monoamines directly target neuronal mitochondrial biogenesis and bioenergetics, and the underlying mechanisms. The focus of our studies has been to address the impact of 5-HT on mitochondrial biogenesis and function, however it is important to highlight that mitochondria are highly dynamic organelles that constantly undergo fission and fusion. We have not assessed effects of 5-HT on mitochondrial morphology and dynamics, which merit future investigation.

The dose-dependent effects of 5-HT on mitochondrial biogenesis and bioenergetics were observed in the micromolar concentration range. 5-HT is reported to exhibit both synaptic and extrasynaptic effects, and while synaptic vesicular concentrations are estimated as being as high as 6 mM, 5-HT can diffuse > 20μm away to extrasynaptic sites, where concentrations fall into the micromolar and nanomolar range (37–42). The doses of 5-HT used in our study fall within this physiologically relevant range, and have been used in several previous studies (37, 38, 43–45). Pharmacological and genetic loss of function studies, clearly demonstrate that the effects of 5-HT on mtDNA content, *Ppargc1a* and *Sirt1* expression and ATP levels are mediated via the 5-HT_2A_ receptor. This is corroborated by genetic rescue of 5-HT_2A_ receptor expression in cortical neurons derived from 5-HT_2A_^-/-^ mice, in which the mitochondrial effects of 5-HT are completely restored. Strikingly, while these effects of 5-HT on mtDNA and ATP levels are completely blocked by both pharmacological blockade or genetic loss of the 5-HT_2A_ receptor, we do not observe a basal change in these measures. This indicates that while the 5-HT_2A_ receptor is essential to mediate the mitochondrial effects of 5-HT, the absence of baseline mitochondrial changes suggests functional redundancy in the pathways that modulate neuronal mitochondrial biogenesis and function. The effects of 5-HT on mtDNA, gene expression, ATP levels and OXPHOS are phenocopied by 5-HT_2A_ receptor stimulation of cortical neurons, and involve the PLC and MAPK, but not the PI3K-Akt, signalling pathways (Fig. 6). However, we do not preclude a role for additional signaling pathways that could be activated by the 5-HT_2A_ receptor, as we restricted our analysis to only the major signaling cascades reported to lie downstream of 5-HT_2A_ receptor stimulation in cortical neurons (46, 47). Our observations indicate that the 5-HT_2A_ receptor is central to mediating the mitochondrial effects of 5-HT on cortical neurons. It is noteworthy that both hallucinogenic (DOI) and non-hallucinogenic (Lisuride) ligands of the 5-HT_2A_ receptor (48) can enhance mitochondrial biogenesis and ATP levels, suggesting a central role for the 5-HT_2A_ receptor in neuronal bioenergetics. Prior reports indicate that 5-HT_2A_ receptor stimulation enhances neurite outgrowth in cortical neurons (49, 50), and is upstream of the regulation of trophic factors, such as BDNF (51). BDNF has been shown to regulate neuronal mitochondrial biogenesis via PGC-1α, and to enhance mitochondrial docking at synapses (52, 53). This motivates future experiments to examine the possibility of a coordinated interplay between 5-HT, the 5-HT_2A_ receptor and BDNF in regulation of trophic and neuroprotective effects, driven via an influence on mitochondria.

The influence of 5-HT on mitochondria in cortical neurons was initially noted at the level of transcriptional modulation of master regulators of mitochondrial biogenesis (SIRT1, PGC-1α) within 48 h post 5-HT treatment, preceding the effects of enhanced mitochondrial mass and function. This timeline indicates that 5-HT exerts transcriptional control of the SIRT1-PGC-1α axis, with robust increases noted in both mRNA and protein levels, and indicates that 5-HT priming of the SIRT1-PGC-1α axis serves as the driver of the mitochondrial effects of 5-HT. 5-HT_2A_ receptor stimulation by DOI, mimics the effects of 5-HT on *Sirt1* and *Ppargc1a* expression, and the 5-HT mediated transcriptional regulation of *Sirt1* and *Ppargc1a* is blocked by MDL100,907, demonstrating that 5-HT modulates the SIRT1-PGC-1α axis through the 5-HT_2A_ receptor. Our results also underscore the importance of SIRT1 for the effects of 5-HT and DOI on mitochondrial biogenesis and ATP levels. Pharmacological blockade of SIRT1 catalytic activity by EX-527 abrogated the effects of 5-HT on mitochondrial mass and cellular ATP levels. This was supported by studies in cortical cultures derived from Sirt1cKO mice, which indicated that 5-HT failed to evoke changes in ATP levels in the absence of SIRT1. *In vivo* studies indicated that the effects of DOI on mtDNA, ATP levels and expression of *Ppargc1a*, *Tfam*, and *Cycs* in the neocortex, were absent in Sirt1cKO mice. While both pharmacological and genetic loss of function studies revealed a vital role for SIRT1 in mediating the effects of 5-HT/5-HT_2A_ receptor stimulation on mitochondrial biogenesis and function, SIRT1 pharmacological inhibition or genetic loss did not result in any baseline change in mtDNA, ATP levels or in *Ppargc1a*, *Tfam*, *Nrf1*,and *Cycs* expression. In keeping with this observation, prior *in vivo* and *in vitro* studies with pharmacological inhibition or genetic loss of *Sirt1* in mouse embryonic fibroblasts, liver or muscle indicate an absence of baseline changes in multiple mitochondrial parameters, namely ATP and mtDNA levels, gene expression of regulators of mitochondrial biogenesis and function, and ETC complex activity (54–57). This serves to highlight that baseline regulation of mitochondrial biogenesis is likely modulated by multiple pathways. Studies with genetic loss of PGC-1α, have also been similarly suggestive of functional redundancy given they do not show major baseline defects in mitochondrial biogenesis and function (58).

The SIRT1-PGC-1α axis activates gene regulatory programs that endow a cell with the ability to meet energy demands, via modulation of mitochondrial biogenesis, cellular respiration, and energy expenditure (14, 18, 19). SIRT1-mediated deacetylation modulates the transcriptional activity of PGC-1α, a transcriptional co-activator that is the master regulator of mitochondrial biogenesis (14, 18). Mitochondrial biogenesis is a long-term adaptive response, that involves the initiation of a coordinated transcriptional program brought about by the SIRT1-PGC-1α axis based recruitment of NRF-1 and TFAM (14, 17, 19) (Fig. 6). NRF-1 enhances expression of OXPHOS machinery, and promotes the expression of the mitochondrial transcription factor, TFAM, that drives transcription and replication of mtDNA (14, 59). In the context of neurons, the upstream factors that integrate environmental cues and then impinge on the SIRT1-PGC-1α axis to initiate mitochondrial biogenesis and facilitate adaptation in response to altering energetic demands remain elusive. Our findings demonstrate a hitherto unidentified relationship between 5-HT and SIRT1, providing novel evidence that 5-HT is an upstream regulator of SIRT1. A previous study indicates SIRT1 can modulate serotonergic neurotransmission via transcriptional effects on monoamine oxidase A (MAO-A) expression (60), and our findings raise the tantalizing possibility of a reciprocal relationship between 5-HT and SIRT1 in the brain. Given the role of 5-HT in facilitating stress adaptation, this suggests the possibility that 5-HT could serve as a vital intermediary in enhancing stress adaptation of neurons through recruitment of the SIRT1-PGC-1α axis to enhance mitochondrial biogenesis and function, thus endowing neurons with enhanced capacity to buffer stress.

Strikingly, 5-HT resulted in a robust reduction in cellular ROS levels and significantly attenuated the enhanced ROS in cortical neurons subjected to excitotoxic and oxidative stress. The concomitant increases evoked by 5-HT in the ROS scavenging enzymes *Sod2* and *Cat*, suggest a role for these antioxidant enzymes in the effects of 5-HT on ROS. 5-HT/5-HT_2A_ receptor stimulation enhanced cortical neuron viability across a wide-range of doses for kainate and H_2_O_2_, a neuroprotective effect that required SIRT1. Neurons face unique energetic demands and the ability of mitochondria to effectively respond to alterations of environment and buffer the “allostatic” load of stress defines the trajectory for neuronal survival (9, 61, 62). In this context, the robust effects of 5-HT/5-HT_2A_ receptor stimulation on spare respiratory capacity may be of particular relevance. Spare respiratory capacity, through the ability to increase mitochondrial respiration when challenged with enhanced energy demands, equips cells with the ability to buffer extreme stress, and serves as a critical factor determining neuronal survival (63). 5-HT via the 5-HT_2A_ receptor results in increased reserve respiratory capacity, and may contribute to the improved neuronal survival observed in 5-HT and DOI pre-treated neurons when faced with stress. A recent report indicates that 5-HT_1F_ receptor stimulation ameliorates dopaminergic neuronal cell death in an animal model of Parkinson’s disease (21). These findings provide support for the speculative possibility that 5-HT could exert circuit-specific neuroprotective effects via distinct 5-HT receptors, promoting neuronal survival through effects on mitochondrial biogenesis, respiration and function. Thus far there remains a limited understanding of potential drug targets to induce mitochondrial biogenesis and modulate mitochondrial function (64–67). Given an emerging role for mitochondrial biogenesis and function as a putative therapeutic target in neuropsychiatric and neurodegenerative disorders (68–71), our work supports the notion that the 5-HT_2A_ receptor may serve as putative drug target to enhance mitochondrial biogenesis and function.

In conclusion, our findings demonstrate that 5-HT can increase mitochondrial biogenesis and function in cortical neurons, via a 5-HT_2A_ receptor-dependent recruitment of the SIRT1-PGC-1α axis. 5-HT, through the 5-HT_2A_ receptor, exerts robust neuroprotective effects on cortical neurons subjected to neurotoxic insults. SIRT1 lies downstream of 5-HT, through the 5-HT_2A_ receptor in cortical neurons, and is essential for the effects of 5-HT on mitochondrial biogenesis, gene expression of regulators of mitochondrial biogenesis, ATP levels and enhanced cell viability under excitotoxic and oxidative stress. These mitochondrial effects of 5-HT bear significance, in relation to the influence of 5-HT, in promoting cell survival, neuronal plasticity, stress adaptation and regulation of senescence/aging.

## Materials and Methods

### Animals

Timed pregnant Sprague-Dawley dams were bred in the Tata Institute of Fundamental Research (TIFR) animal facility and embryos derived at embryonic day 18.5 (E18.5) were used for all cortical culture experiments, with the exception of experiments with cortical cultures established from specific mouse mutant lines. Mouse cortical cultures were generated from E18.5 embryos derived from wild-type and serotonin_2A_ receptor knockout (5-HT_2A_^-/-^) dams, with the possibility of Cre-mediated rescue of 5-HT_2A_ receptor expression (48). Mouse cortical cultures were also generated from embryos derived from bigenic E18.5 Emx1-Cre;Sirt1^+/+^ (wild-type: WT) and Emx1-Cre;Sirt1^-/-^ (Sirt1cKO) dams (Emx1-Cre: Jax-mice-ID 005628) (SirT1^co^: Jax-mice-ID 008041) (72). For *in vivo* experiments, male Sprague Dawley rats (5-6 months), WT and Sirt1cKO mice (15 months) were used. All animals were group housed, maintained on a 12 h light–dark cycle with *ad libitum* access to food and water. Experimental procedures were in accordance with the guidelines of the Committee for Supervision and Care of Experimental Animals (CPCSEA), Government of India, and were approved by the TIFR Institutional Animal Ethics committee (CPCSEA-56/1999).

### Primary cortical culture

Cortices were dissected from E18.5 rat or mouse embryos and were cultured in Neurobasal medium supplemented with 2% B27 and 0.5 mM L-glutamine (Thermo Fisher Scientific, USA). For specific sets of experiments cortical cultures generated from 5-HT_2A_^-/-^mouse embryos were transduced with rAAV8-CaMKIIα-GFP-Cre (UNC Vector Core facility, NC, USA) on DIV 2 to restore *Htr2a* expression (5-HT_2A_^-/-Res^).

### Drug treatment paradigms

For dose response studies, rat cortical neurons were treated with 5-HT (10, 50 and 100 μM) for six days. For mitochondrial marker analysis, mitotracker imaging experiments and reactive oxygen species (ROS) measurements, rat cortical cultures were treated with 5-HT (100 μM) for six days. For time-point analysis, cortical neurons were treated with 5-HT (100 μM) for durations of 24 h, 48 h, 72 h, and 6 days. Cortical cultures were also treated with norepinephrine or dopamine (10, 100 μM) for six days to assess influence of other monoamines. Mouse cortical cultures from 5-HT_2A_^-/-^, 5-HT_2A_^-/-Res^, SIRT1cKO and their respective controls were treated with 5-HT (100 μM) for six days. In experiments with 5-HT receptor antagonists, specific inhibitors of kinase pathways or the SIRT1 inhibitor, rat cortical neurons were treated with 5-HT (100 μM) for 72 h in the presence of the 5-HT_2A_ receptor antagonist, MDL100,907 (10 μM), the 5-HT_1A_ receptor antagonist, WAY100,635 (10 μM), MEK inhibitor U0126 (10 μM), PLC inhibitor U73122 (5 μM), PI3-kinase inhibitor LY294002 (10 μM) or SIRT1 inhibitor EX-527 (10 μM). For experiments with the 5-HT_2A_ receptor agonist, rat cortical neurons were treated with DOI (5, 10 μM) for six days or Lisuride (10 μM) for 72 h. For mitochondrial respiration analysis, cortical neurons were seeded at an equal density of 10 × 10^4^ cells per well of the Seahorse XF24 cell culture microplate (Seahorse Bioscience, Agilent Technologies, CA, USA), and treated with 5-HT (100 μM) or DOI (10 μM) for 72 h. For studies addressing the influence of 5-HT in protection against excitotoxic or oxidative stress, cell viability and cellular reactive oxygen species (ROS) were assessed in cortical neuron cultures pretreated with 5-HT (50, 100 μM, 6 days) followed by challenge with kainate (100, 200 μM; 3.5 h) or H_2_O_2_ (100, 200 μM; 7 h). The contribution of SIRT1 to the effects of 5-HT on neuronal viability following kainate (10 to 1000 μM) or H_2_O_2_ (10 to 1000 μM) treatment was determined using (1) the SIRT1 inhibitor EX-527 (10 μM), or (2) cortical neurons derived from Sirt1cKO and WT mice. The influence of DOI (10 μM) in neuroprotection against kainate (100, 200 μM; 3.5 h) or H_2_O_2_ (100, 200 μM; 7 h) was determined by assessing cell viability and the contribution of SIRT1 in these effects of DOI was determined using EX-527 treatment (10 μM). Controls involved treatment of cultures with vehicle (water), with the exception of U73122, U0126, LY294002, EX-527, Lisuride and DOI where the vehicle was 0.1% DMSO. For *in vivo* experiments, Sprague-Dawley rats, Sirt1cKO and WT mice received intraperitoneal administration of DOI (2 mg/kg) or vehicle (0.9 % saline) once daily for four days and were sacrificed two hours after the last treatment.

### Quantitative real time polymerase chain reaction

RNA was reverse transcribed to cDNA and amplified using gene specific primers as detailed in supplementary methods.

### Mitochondrial DNA (mtDNA) levels

Mitochondrial DNA (mtDNA) levels were determined by quantitative real time PCR (qPCR). Levels of cytochrome B - a mitochondrial genome encoded gene were normalised to levels of a nuclear encoded gene cytochrome C and quantified by the ΔΔCt method.

### Immunofluorescence

Immunofluorescence experiments involved staining with primary and secondary antibodies prior to visualization on the Olympus Fluoview 1000 confocal laser scanning microscope. Intensity of VDAC staining per micron neurite length was quantitated using ImageJ software (NIH, USA) in 15-20 neurons per treatment condition per experiment.

### Western blot analysis

Cell lysates were resolved by SDS-PAGE, followed by Western blotting and probed with primary and secondary antibodies, as detailed in supplementary methods.

### Mitotracker Staining

Cortical neurons were loaded with mitotracker green (20 nM, Invitrogen) for 45 min, and images were recorded using an Olympus Fluoview 1000 confocal laser scanning microscope.

The intensity of staining in neurites was quantitated over a distance of 10 μm commencing from the point proximal to the cell body using ImageJ^®^ software (NIH, USA) in 15-20 neurons per treatment condition per experiment.

### Cellular ATP assay

Cellular ATP levels were determined using a commercial ATP bioluminescent assay kit (Sigma-Aldrich) and normalized to total protein content per sample, and expressed as fold change of control.

### Mitochondrial Respiration Analysis

Mitochondrial respiration measured as oxygen consumption rate (OCR) was determined in cortical neurons using the Seahorse XFe24 extracellular flux analyzer (Seahorse Bioscience, Agilent Technologies, CA, USA), with the commercially available XF Cell Mito Stress Test Kit (Seahorse Bioscience), as per the manufacturer’s instructions. OCR (pmol min^−1^) was measured at baseline and in the presence of the ATP synthase inhibitor, oligomycin (1 μM), uncoupling agent FCCP (2 μM) and ETC complex I inhibitor rotenone (0.5 μM) to determine basal respiration, ATP coupled respiration, maximal respiration, non-mitochondrial respiration and spare respiratory capacity.

### Cellular ROS measurement

Cortical neurons were incubated with carboxy-H_2_DCFDA (25 μM, Molecular Probes) and ROS levels were assessed by fluorescence measurements (Ex/Em: 495/529 nm) with representative images captured using the Olympus Fluoview 1000 confocal laser scanning microscope.

### Statistical Analysis

Data were subjected to statistical analysis using GraphPad Prism 7 and GraphPad InStat (GraphPad Software Inc, USA). Significance between two groups was evaluated using the unpaired Student’s *t*-test. Experiments with multiple group comparisons were subjected to one-way, two-way or three-way ANOVA analysis followed by *post-hoc* Tukey comparisons. Statistical significance was set at *p* < 0.05.

## Acknowledgements

This research was supported by TIFR intramural funds (U.K. & V.A.V). We acknowledge Dr. Suryavanshi of the TIFR animal facility for technical assistance.

## Author Contributions

SF, ABV, UK and VAV designed research; SF, SD, BK and SG performed research; NW and JG contributed new reagents/analytic tools; SF, SD, ABV, UK and VAV analysed data; SF, UK and VAV wrote the paper; UK and VAV supervised the project.

Authors declare no conflict of interest.

## Supplementary Information

### Supplementary Methods

#### Primary cortical culture

Dams were euthanized with CO_2_ and embryos were collected in ice cold Ca^2+^ and Mg^2+^ free Hanks’ balanced salt solution (HBSS) supplemented with 300 mM HEPES. Cortices were dissected in ice cold minimum essential medium, enzymatically dissociated in 0.05% trypsin/EDTA for 10 min, and triturated to obtain a single cell suspension in Neurobasal medium supplemented with 2% B27 and 0.5 mM L-glutamine. Cells were seeded in plates coated with poly-D-lysine (0.1 mg/ml, Sigma-Aldrich, USA) at a density of 1 million cells per 9.6 cm^2^. Neurons were cultured at 37°C with 5% CO_2_ and 95% humidity with a half-medium change every 2 days. Cortical neurons were allowed to differentiate for at least seven days *in vitro* (DIV 7) prior to treatment (1, 2). All reagents were purchased from Thermo Fisher Scientific, (USA) unless specified.

#### Drugs

All compounds were purchased from Tocris Bioscience (United Kingdom), except 5-HT, DOI, norepinephrine, WAY100,635, EX-527, H_2_O_2_ and kainate (Sigma-Aldrich, USA). The concentration of neurotransmitters used was based on prior studies (3–8).

#### Quantitative real time polymerase chain reaction

RNA was extracted from cortical neurons using the commercially available RNeasy Micro/Mini kit (Qiagen). 50 ng of RNA per sample was reverse transcribed to cDNA using random hexamers and the Superscript IV reverse transcription kit (Invitrogen, USA). Quantitative real time PCR was performed using gene specific primers and cDNA was amplified in a Light Cycler 96 (Roche Applied Science, Switzerland) real time PCR system using KAPA SYBR^®^ FAST Universal 2X qPCR Master Mix (Kapa Biosystems). Gene expression levels were normalized to the endogenous 18S ribosomal RNA per sample, and the relative expression levels between control and treated samples were computed by the ΔΔCt method, as described previously (9). Data are represented as fold change ± SEM as compared to control.

#### Mitochondrial DNA levels

Mitochondrial DNA (mtDNA) levels were compared in control versus treated cortical neurons by quantitative real time PCR. Total DNA was extracted from cells using the commercially available All Prep DNA/RNA Mini kit (Qiagen, Germany). Levels of cytochrome B - a mitochondrial genome encoded gene were normalized to levels of a nuclear encoded gene cytochrome C. Relative mitochondrial DNA levels between groups were quantified by the qqCt method as described previously (10).

#### Immunofluorescence

For immunostaining, cortical neurons were fixed in 4% paraformaldehyde, permeabilized with 0.2% Triton X-100, followed by blocking in 10% horse serum. Cells were then incubated with primary antibodies, which included the rabbit anti-VDAC (1:200, Abcam, MA, USA) or goat anti-5-HT_2A_ (1:250, Santacruz Biotechnologies, CA, USA), incubated along with the pan neuronal marker mouse anti-MAP2 (1:500, Sigma-Aldrich) overnight at 4°C. Cells were washed and incubated with secondary antibodies, Alexa Fluor 488 conjugated anti-rabbit (1:250, Molecular probes) or Alexa Fluor 488 conjugated anti-goat (1:250, Molecular probes, CA, USA) or Alexa Fluor 568 conjugated anti-mouse (1:250, Molecular probes, CA, USA) respectively for 2 hours, followed by washes. The nuclei were counterstained using Hoechst, mounted in Vectashield (Vector laboratories, CA, USA) and images were captured on the Olympus Fluoview 1000 confocal laser scanning microscope.

#### Western blot analysis

Control and treated cortical neuron cultures were lysed in ice cold Radioimmunoprecipitat—ion assay (RIPA) buffer (10 mM Tris-Cl (pH 8.0), 1 mM EDTA, 0.5 mM EGTA, 1% NP-40, 0.1% sodium deoxycholate, 0.1% SDS, 140 mM NaCl), supplemented with protease and phosphatase inhibitors (Roche Applied Science). Lysates were centrifuged at 12000 rpm for 30 min and supernatant was collected. Protein estimation was performed using the QuantiPro BCA (Bicinchoninic Acid) assay kit (Sigma-Aldrich) and lysates were resolved by sodium dodecyl sulfate polyacrylamide gel electrophoresis (SDS-PAGE). Proteins were transferred onto polyvinylidene fluoride (PVDF, Merck Millipore, MA, USA) membranes, blocked in 5% milk in Tris Buffered Saline-Tween (0.1%) and probed with primary antibodies in 5% BSA (bovine serum albumin) overnight at 4°C. Primary antibodies included mouse anti-PGC-1α (1:300, Calbiochem), rabbit anti-TFAM (1:1000, Sigma-Aldrich), rabbit anti-VDAC (1:250, Abcam), mouse anti-Cyt C (1:250, Abcam), mouse anti-SIRT1 (1:250, Sigma-Aldrich), mouse anti-actin (1:4000, Sigma-Aldrich), rabbit anti-βIII-tubulin (1:1000), rabbit anti-phospholipase C (PLC) beta 3 (1:1000), rabbit anti-phospho PLC beta 3 (1:1000), rabbit anti-Akt (1:2000), rabbit anti-phospho Akt (1:1000), rabbit anti-p44/42 MAPK (ERK1/2) (1:1000) and rabbit anti-phospho p44/42 MAPK (ERK1/2) (1:500) (Cell Signaling Technology, MA, USA). Blots were washed and incubated with goat anti-rabbit IgG peroxidase labelled or rabbit anti-mouse IgG peroxidase labelled (1:7000, Sigma-Aldrich) secondary antibodies for 2 hours at room temperature. Signal was detected using a chemiluminescence kit (Thermo Fisher Scientific), and bands were visualized on x-ray films (Fiji films) or the GE Amersham Imager 600. Actin or tubulin were used as loading controls for normalization as indicated, and the relative density of bands was quantitated using ImageJ software (NIH, USA).

#### Cellular ATP

Cortical neurons were lysed in boiling water and lysate was centrifuged at 12,000 rpm for 20 min at 4°C. The ATP levels in the supernatant were quantified using the ATP bioluminescent assay kit (Sigma-Aldrich), by mixing the luciferin substrate and luciferase enzyme mix with equal amounts of supernatant in a 96 well plate. The light emitted is proportional to the ATP consumed in the reaction, and was measured using a luminometer (Berthold Technologies, Germany). ATP levels were normalized to protein content of each sample, estimated using a BCA protein assay kit (Sigma-Aldrich) and expressed as fold change of treated over control.

#### Mitochondrial Respiration Analysis

Cortical neurons were seeded at an equal density of 10 × 10^4^ cells per well of the Seahorse XF24 cell culture microplate and differentiated in Neurobasal medium. Cells were treated with 5-HT or DOI for 72 h, with 4-5 replicate wells per group. On the day of the assay, control and treated neurons were washed twice in assay medium, and 500 μL of assay medium pre-warmed to 37°C, was added in each well. Assay medium comprised of Seahorse XF base medium supplemented with 25 mM glucose (Sigma), 1 mM pyruvate (Sigma), 2 mM glutamine (Thermo Fisher Scientific) pH 7.4. 2% B27 (Thermo Fisher Scientific) was added prior to the addition of the medium to the wells. The plate was equilibrated in a non–CO_2_ incubator for 45 min at 37°C, and thereafter loaded into the Seahorse XF24 analyzer. Oxygen Consumption rate (OCR) (pmol min^−1^) was measured baseline in assay medium, and thereafter cells were exposed to sequential injections of oligomycin (1 μM, ATP synthase complex V inhibitor), FCCP (2 μM, collapses the proton gradient, disrupts the mitochondrial membrane potential and uncouples oxygen consumption from ATP production) and rotenone (0.5 μM, complex I inhibitor), with three consecutive OCR measurements taken after each injection. OCR measurements at baseline and following the three serial injections, were used to calculate basal respiration, ATP production, maximal respiration, non-mitochondrial respiration and spare respiratory capacity. OCR measurements were normalized to blank wells and cell count in each well, and expressed as pmol/min/10^4^ cells. Experiments were repeated thrice and values from all the repeats were collated when calculating mitochondrial parameters. A representative trace of OCR per treatment condition is depicted.

#### Cellular ROS Measurement

To assess levels of cellular reactive oxygen species (ROS), carboxy-H_2_DCFDA (25 μM) was added to the medium for 30 min at 37°C. Cells were then washed twice and images were captured using the Olympus Fluoview 1000 confocal laser scanning microscope. Alternatively, to quantitate the fluorescence, cells were lysed in RIPA buffer and fluorescence was read per treatment per well at excitation/emission of 495/529nm, and normalized to protein content per well. Protein was estimated by BCA method (Sigma).

#### Data Availability Statement

Detailed protocol, raw data and materials will be made available upon request. This manuscript does not contain any omics datasets.

### Supplementary Figures

**Figure S1-Supplementary data for Figure 1:**
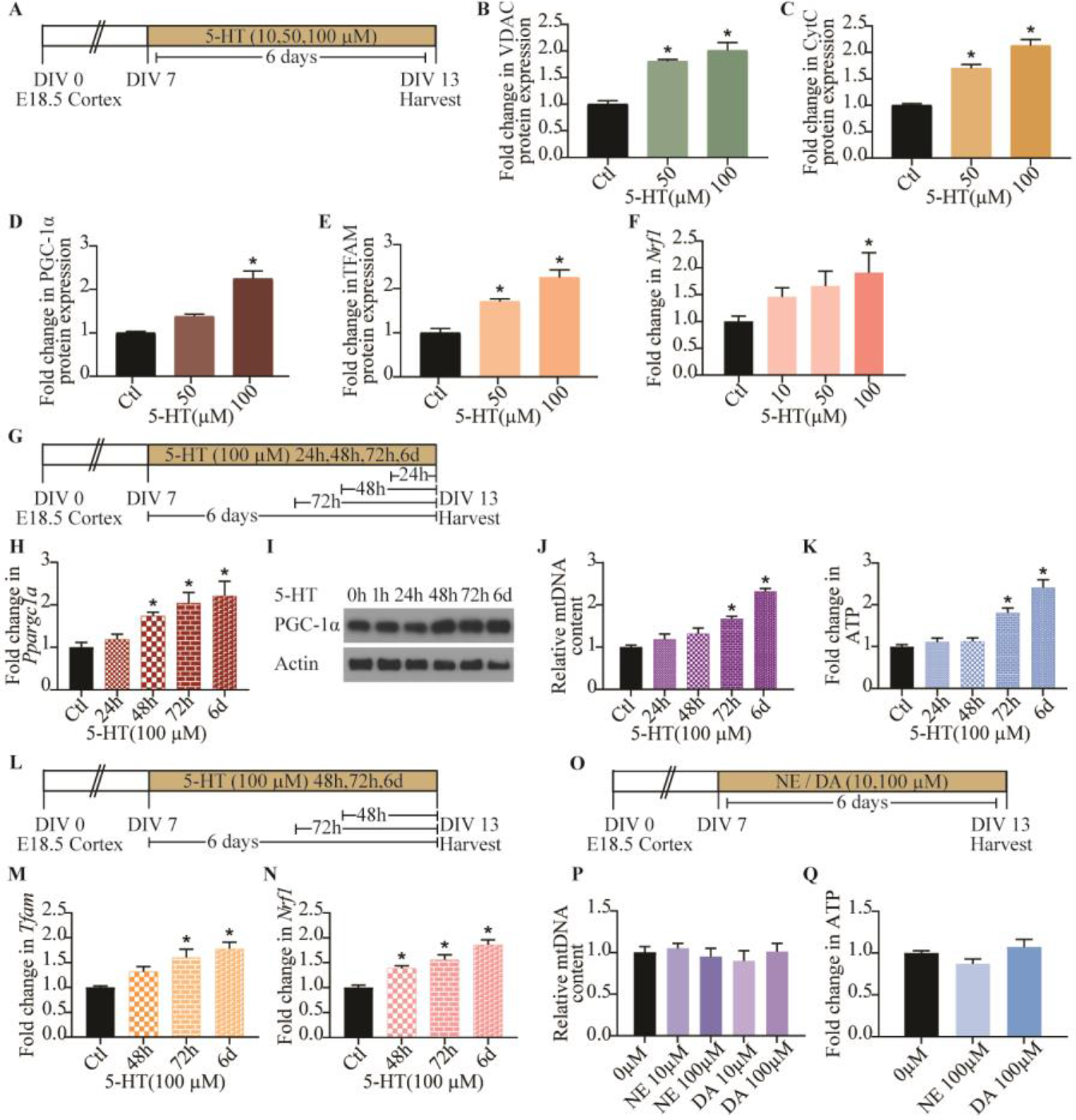
Serotonin regulates mitochondrial biogenesis and function. (A)Shown is a schematic depicting treatment of neurons with increasing doses of 5-HT (10, 50, 100 μM) starting at DIV 7. (B-E) Quantitative densitometric analysis of VDAC (B), Cyt C (C), PGC-1α (D) and TFAM (E) immunoblots from control (Ctl) and 5-HT treated cortical neuron cultures. Results are expressed as fold change of control ± SEM (Compiled results from n = 5-11 per treatment group/N = 2-3, \**p* < 0.05 as compared to control, one-way ANOVA, Tukey’s *post-hoc* test). (F) Quantitative qPCR analysis of *Nrf1* expression from control and 5-HT treated cortical neuron cultures, results are represented as fold change of control ± SEM. (Representative results from n = 4 per treatment group/N = 2, \**p* < 0.05 as compared to control, one-way ANOVA, Tukey’s *post-hoc* test). (G) Shown is a schematic depicting 5-HT (100 μM) treatment of cortical neurons for 24 h, 48 h, 72 h or 6 days and lysed synchronously at DIV 13. (H) qPCR analysis of *Ppargc1a* expression levels in the 5-HT time-course experiment, results are represented as fold change of control ± SEM. (Representative results from n = 4 per treatment group/N = 2, \**p* < 0.05 as compared to control, one-way ANOVA, Tukey’s *post-hoc* test). (I) Shown is a representative immunoblot for PGC-1α with actin as the loading control in the 5-HT time-course experiment at indicated time-points. (J and K) Graphs depict quantitation of mtDNA and ATP levels in the 5-HT time-course experiment at indicated time-points, and results are represented as fold change of control ± SEM. (Representative results from n = 4 per treatment group/N = 2, \**p* < 0.05 as compared to control, one-way ANOVA, Tukey’s *post-hoc* test). (L) Shown is a schematic depicting the treatment paradigm of neurons with 5-HT (100 μM) for 48 h, 72 h or 6 days and lysed synchronously at DIV 13. (M and N) qPCR analysis of *Tfam* (M) and *Nrf1* (N) expression levels in the 5-HT time-course experiment at indicated time-points, and results are represented as fold change of control ± SEM. (Representative results from n = 4 per treatment group/N = 2, \**p* < 0.05 as compared to control, one-way ANOVA, Tukey’s *post-hoc* test). (O) Schematic depicting treatment of neurons with different doses of norepinephrine (NE: 10, 100 μM) or dopamine (DA: 10, 100 μM) from DIV 7 to DIV 13. (P and Q) Graphs depict quantitation of relative mtDNA (P) and ATP (Q) levels in neurons treated with NE or DA. Results are expressed as fold change of control ± SEM. (Representative results from n = 4 per treatment group/N = 2, \**p* < 0.05 as compared to control, one-way ANOVA, Tukey’s *post-hoc* test).

**Figure S2-Suplementary data for Figure 2:**
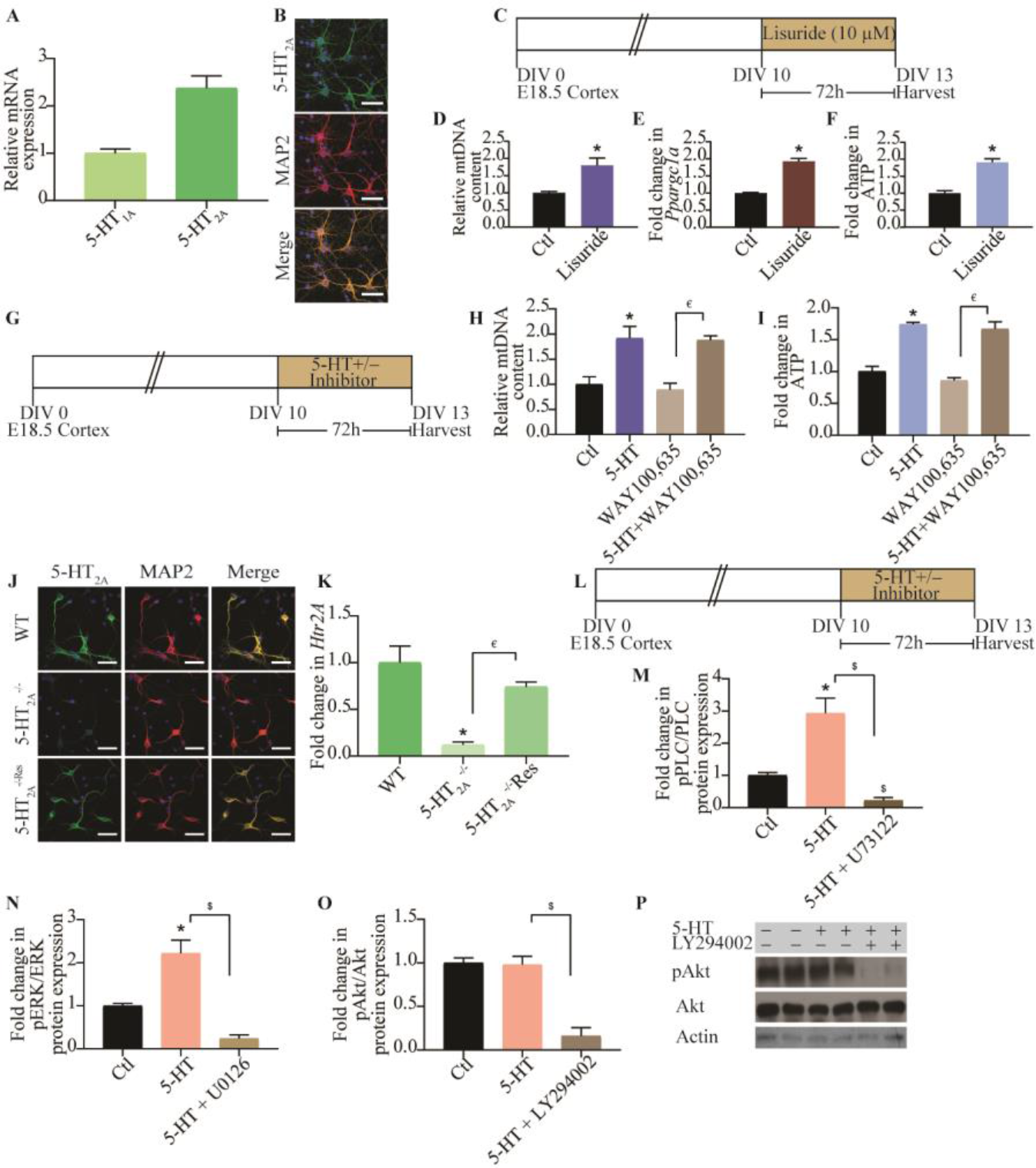
Mitochondrial effects of 5-HT are mediated via the 5-HT_2A_ receptor. Quantitative qPCR analysis of mRNA expression of 5-HT_1A_ and 5-HT_2A_ receptor in cortical neuron cultures. (Representative results from n = 4 per treatment group/N = 2). (B) Shown are representative images for 5-HT_2A_ receptor immunofluorescence (green), neuronal marker MAP2 (red) and merge (yellow) from cortical neuron cultures. Scale bar: 50 μm. Magnification: 60X. (C) Shown is a schematic depicting the treatment paradigm for cortical neuron cultures with Lisuride (10 μM), for 72 h commencing DIV 10. (D-F) Quantitation of mtDNA (D), *Ppargc1a* mRNA (E) and ATP (F) levels in cortical neurons treated with Lisuride are represented as fold change of control (Ctl) ± SEM. (Representative results from n = 4-8 per treatment group/N = 2, \**p* < 0.05 as compared to control, unpaired Student’s *t*-test). (G) Shown is a schematic depicting the treatment paradigm of cortical neuron cultures with 5-HT in the presence or absence of the selective 5-HT_1A_ receptor antagonist, WAY100,635 commencing DIV 10. (H and I) Graphs depict quantitative analysis for mtDNA (H) and ATP (I) levels with results expressed as fold change of control (Ctl) ± SEM. (Representative results from n = 4 per treatment group/N = 2, \**p* < 0.05 as compared to control, ^€^*p* < 0.05 as compared to WAY100,635, two-way ANOVA, Tukey’s *post-hoc* test). (J)Shown are representative images for 5-HT_2A_ receptor immunofluorescence (green), MAP2 (red) and merge (yellow) in WT, 5-HT_2A_^-/-^ and 5-HT_2A_^-/-Res^ cortical neuron cultures. Scale bar: 50 μm. Magnification: 60X. (K) Quantitative analysis of *Htr2A* mRNA levels in cortical neuron cultures from WT, 5-HT_2A_^-/-^ and 5-HT_2A_^-/-Res^ indicate reduced *Htr2A* levels in 5-HT_2A_^-/-^ cultures, with rescue of expression in the 5-HT_2A_ group. Results are expressed as the fold change of WT ± SEM. (Representative results from n = 4 per treatment group/N = 2, \**p* < 0.05 as compared to WT, ^€^*p* < 0.05 as compared to 5-HT_2A_^-/-^, one-way ANOVA, Tukey’s *post-hoc* test). (L) Shown is a schematic depicting the treatment paradigm of cortical neurons with 5-HT (100 μM) in the presence or absence of signaling inhibitors for PLC (U73122, 5 μM), MEK (U0126, 10 μM) and PI3K (LY294002, 10 μM). (M-P) Quantitative densitometric analysis of pPLC/PLC levels (M), pERK/ERK levels (N) and pAkt/Akt (O) levels with ratios normalized to actin or tubulin as loading controls. Results are expressed as fold change of control ± SEM. (Compiled results from n = 4-5 per treatment group/N = 3, \**p* < 0.05 as compared to control, ^$^p < 0.05 as compared to 5-HT treated group, one-way ANOVA, Tukey’s *post-hoc* test). (P) Shown is a representative immunoblot of pAkt and Akt in control and 5-HT treated cells in the presence of absence of the PI3K inhibitor, LY294002 which resulted in a robust inhibition of pAkt levels in 5-HT treated cortical neurons.

**Figure S3-Suplementary data for Figure 3:**
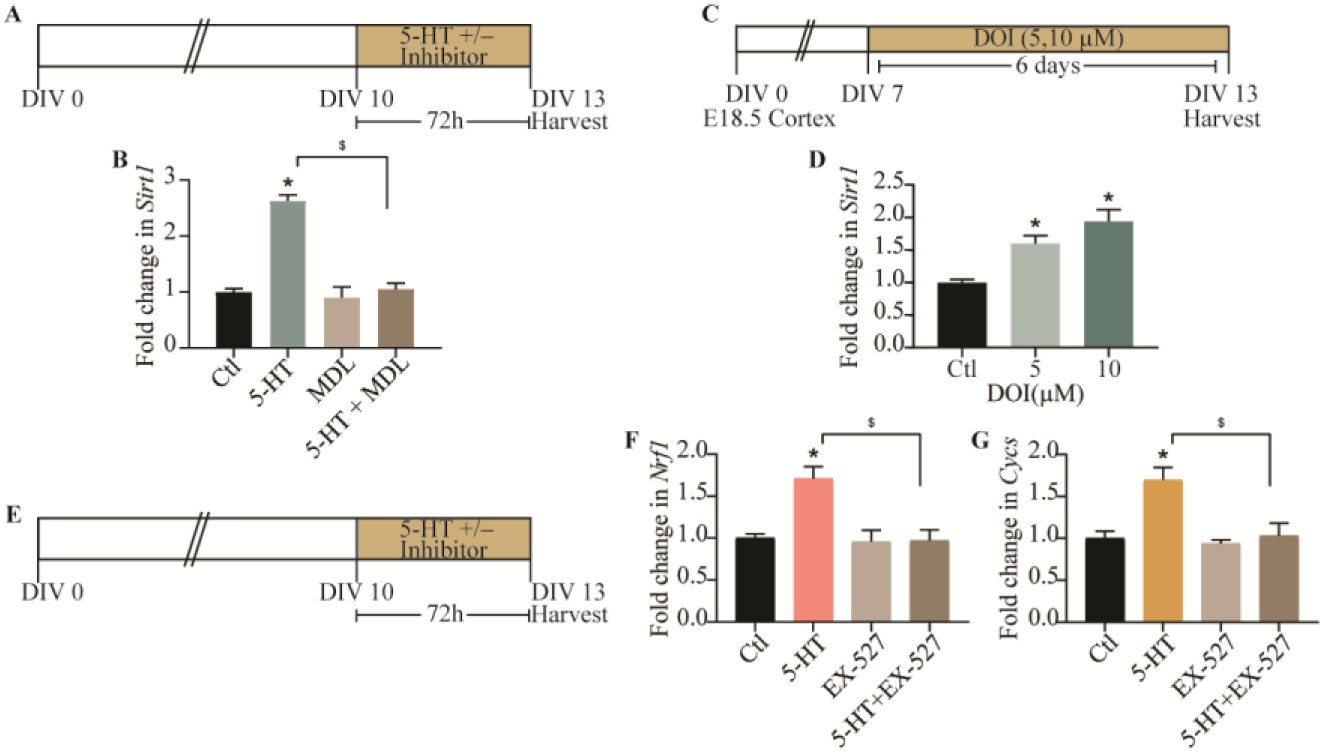
SIRT1 is required for the effects of 5-HT on mitochondria. (A)Shown is a schematic depicting the treatment paradigm of neurons with 5-HT (100 μM) in the presence or absence of the 5-HT_2A_ receptor antagonist MDL100,907 (MDL) (10 μM) commencing DIV 10. (B) Graph depicts quantitation of *Sirt1* mRNA levels in cortical neurons treated with 5-HT in the presence or absence of MDL100,907 (MDL), and expressed as fold change of control ± SEM. (Representative results from n = 4 per treatment group/N = 2, \**p* < 0.05 as compared to control, ^$^*p* < 0.05 as compared to 5-HT treated group, two-way ANOVA, Tukey’s *post-hoc* test). (C) Shown is a schematic depicting the treatment paradigm of neurons with increasing doses of DOI (5, 10 μM) commencing DIV 7. (D) Quantitative qPCR results for *Sirt1* mRNA levels are expressed as fold change of control ± SEM. (Representative results from n = 4 per treatment group/N = 2, \**p* < 0.05 as compared to control, one-way ANOVA, Tukey’s *post-hoc* test). (E) Shown is a schematic depicting the treatment paradigm of neurons with 5-HT (100 μM) in the presence or absence of the Sirt1 inhibitor, EX-527 (10 μM). (F and G) Quantitation of mRNA expression of *Cycs* (F) and *Nrf1* (G) in cortical neurons treated with 5-HT in the presence or absence of EX-527 are expressed as fold change of control ± SEM. (Representative results from n = 4 per treatment group/N = 2, \**p* < 0.05 as compared to control, ^$^*p* < 0.05 as compared to 5-HT treated group, two-way ANOVA, Tukey’s *post-hoc* test).

**Figure S4-Supplementary data for Figure 4:**
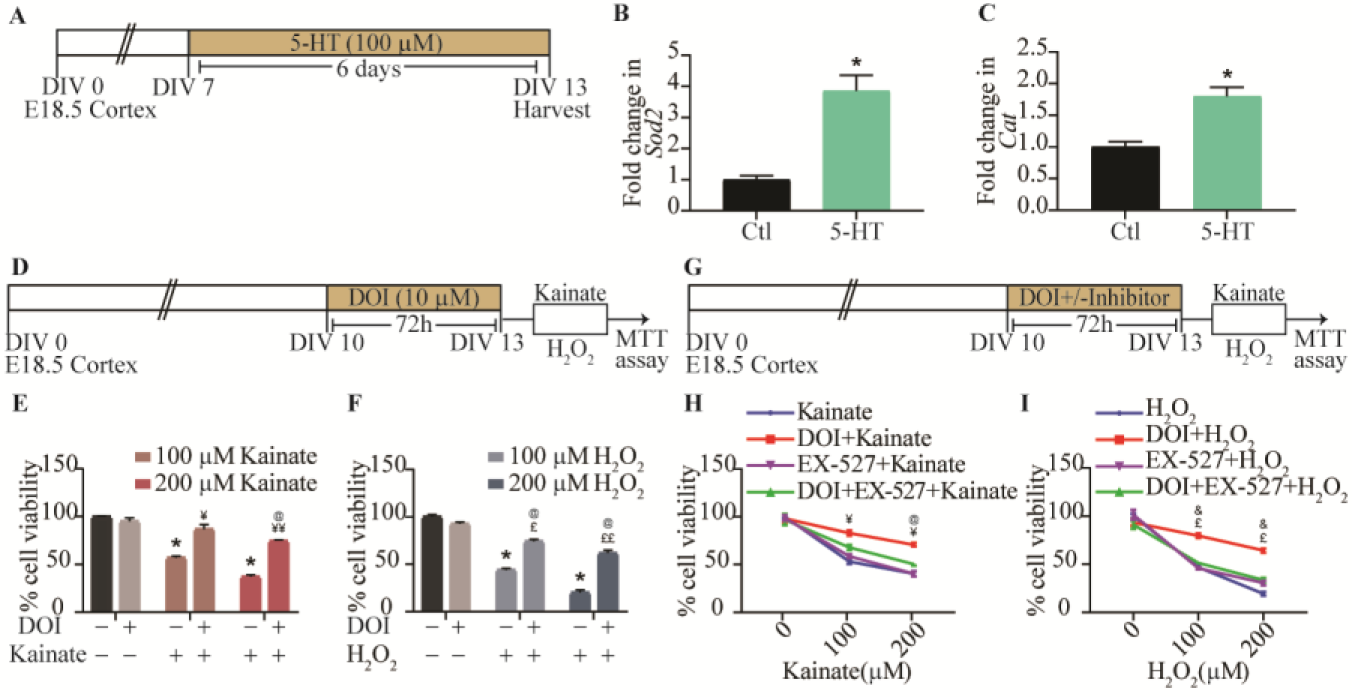
5-HT_2A_ receptor activation exerts neuroprotective effects against excitotoxic and oxidative stress. (A)Shown is a schematic depicting the treatment paradigm of neurons with 5-HT (100 μM) commencing DIV 7. (B and C) qPCR analysis of *Sod2* and *Cat* mRNA levels in 5-HT treated cortical neurons are expressed as fold change of control (Ctl) ± SEM. (Representative results from n = 4 per treatment group/N = 2, \**p* < 0.05 as compared to control, unpaired Student’s *t*-test). (D) Shown is a schematic depicting the treatment paradigm of cortical neurons with the 5-HT_2A_ receptor agonist, DOI (10 μM) commencing DIV 10 followed by a challenge with kainate (100, 200 μM) or H_2_O_2_ (100, 200 μM) on DIV 13 and analysis of cell viability. (E) Graph depicts cell viability assessed by the MTT assay in cortical neurons challenged with kainate (100, 200 μM) with or without pretreatment with DOI (10 μM). Results are expressed as percent of control cell viability ± SEM. (Representative results from n = 3 per treatment group/N = 2, \**p* < 0.05 as compared to control, ^¥^*p* < 0.05 as compared to 100 μM kainate treated group, ^¥¥^*p* < 0.05 as compared to 200 μM kainate treated group, ^@^*p* < 0.05 as compared to DOI treated group, two-way ANOVA, Tukey’s *post-hoc* test). (F) Graph depicts cell viability assessed by the MTT assay in cortical neurons challenged with H_2_O_2_ (100, 200 μM) in cortical neurons with or without pretreatment with DOI (10 μM). Results are expressed as percent of control cell viability ± SEM. (Representative results from n = 3 per treatment group/N = 2, \**p* < 0.05 as compared to control, ^£^*p* < 0.05 as compared to 100 μM H_2_O_2_ treated group, ^££^*p* < 0.05 as compared to 200 μM H_2_O_2_ treated group, ^@^*p* < 0.05 as compared to DOI treated group, two-way ANOVA, Tukey’s *post-hoc* test). (G) Shown is a schematic depicting the treatment paradigm of neurons with DOI (10 μM) in the presence or absence of the SIRT1 inhibitor, EX-527 (10 μM), followed by challenge with kainate or H_2_O_2_ (100, 200 μM) and analysis of cell viability using the MTT assay. (H) Shown is a line graph for cell viability of cortical neurons in response to kainate (100, 200 μM) with treatment groups of kainate (blue), DOI + kainate (red), EX-527 + kainate (purple), DOI + EX-527 + kainate (green) expressed as per cent of control cell viability ± SEM. (Representative results from n = 3 per treatment group/N = 2, ^¥^*p* < 0.05 as compared to kainate treatment group, ^@^*p* < 0.05 as compared to DOI + EX-527 + kainate treated group, three-way ANOVA, Tukey’s *post-hoc* test). (I) Shown is a line graph for cell viability of cortical neurons in response to H_2_O_2_ (100, 200 μM) with treatment groups of H_2_O_2_ (blue), DOI + H_2_O_2_ (red), EX-527 + H_2_O_2_ (purple), DOI + EX-527 + H_2_O_2_ (green) expressed as per cent of control cell viability ± SEM. (Representative results from n = 3 per treatment group/N = 2, ^£^*p* < 0.05 as compared to H_2_O_2_ treated group, ^&^*p* < 0.05 as compared to DOI + EX-527 + H_2_O_2_ treated group, three-way ANOVA, Tukey’s *post-hoc* test).

### List of Primers

**Table S1:**
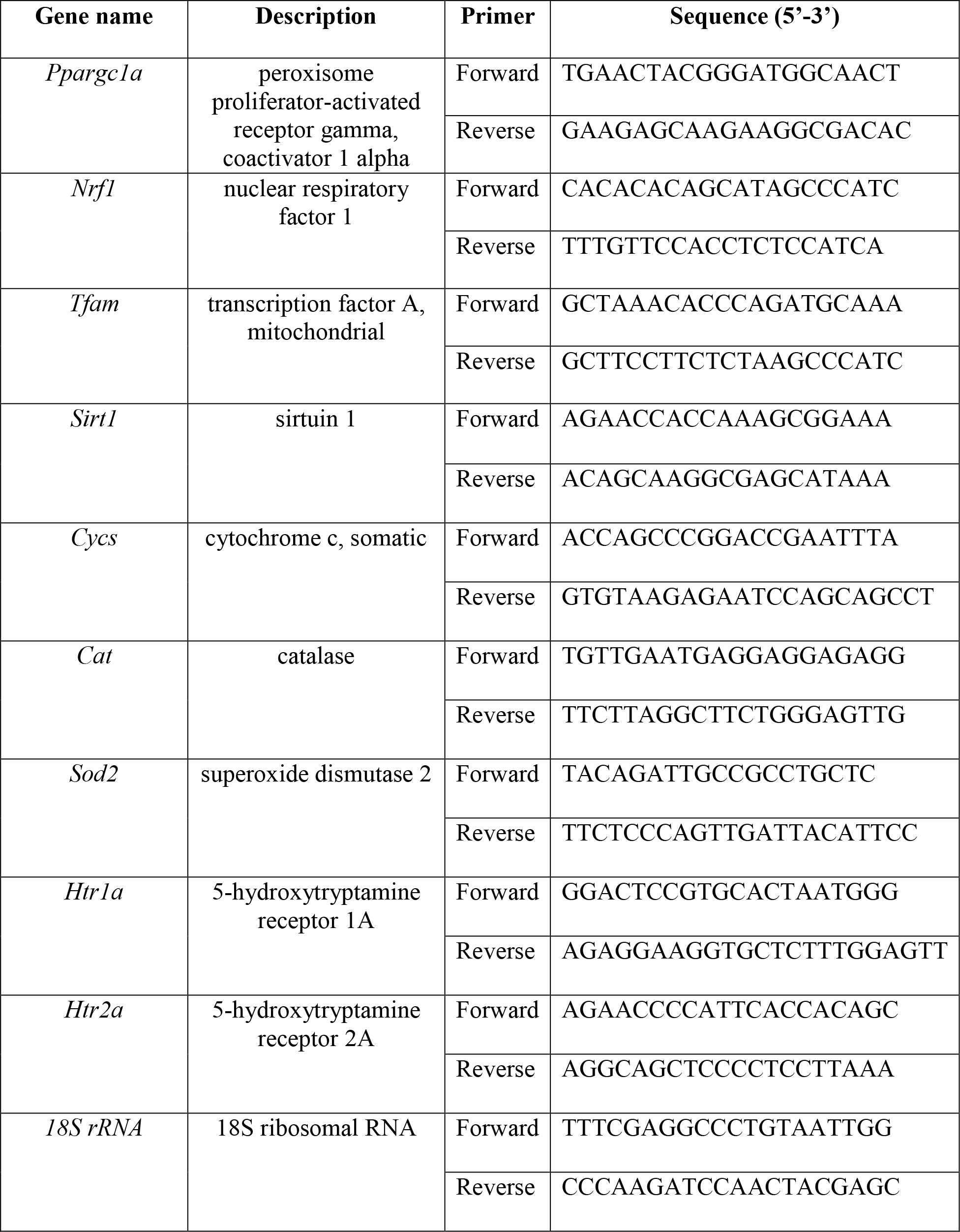
Primer sequences used for quantitative PCR analysis of rat cDNA

**Table S2:**
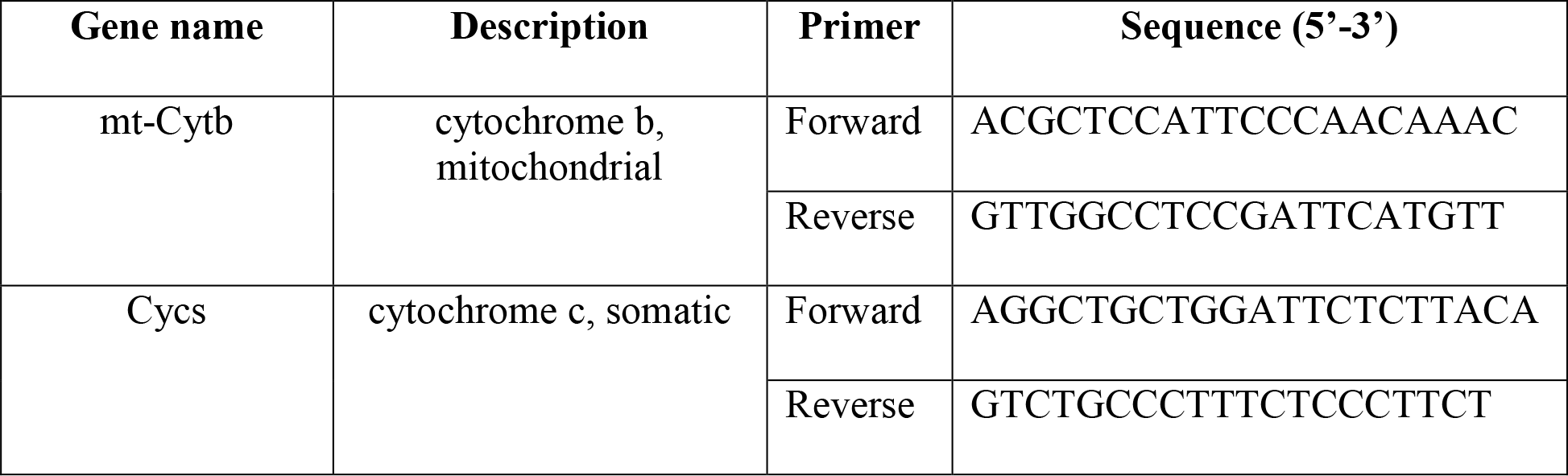
Primer sequences used for relative mitochondrial DNA (mtDNA) analysis in rat

**Table S3:**
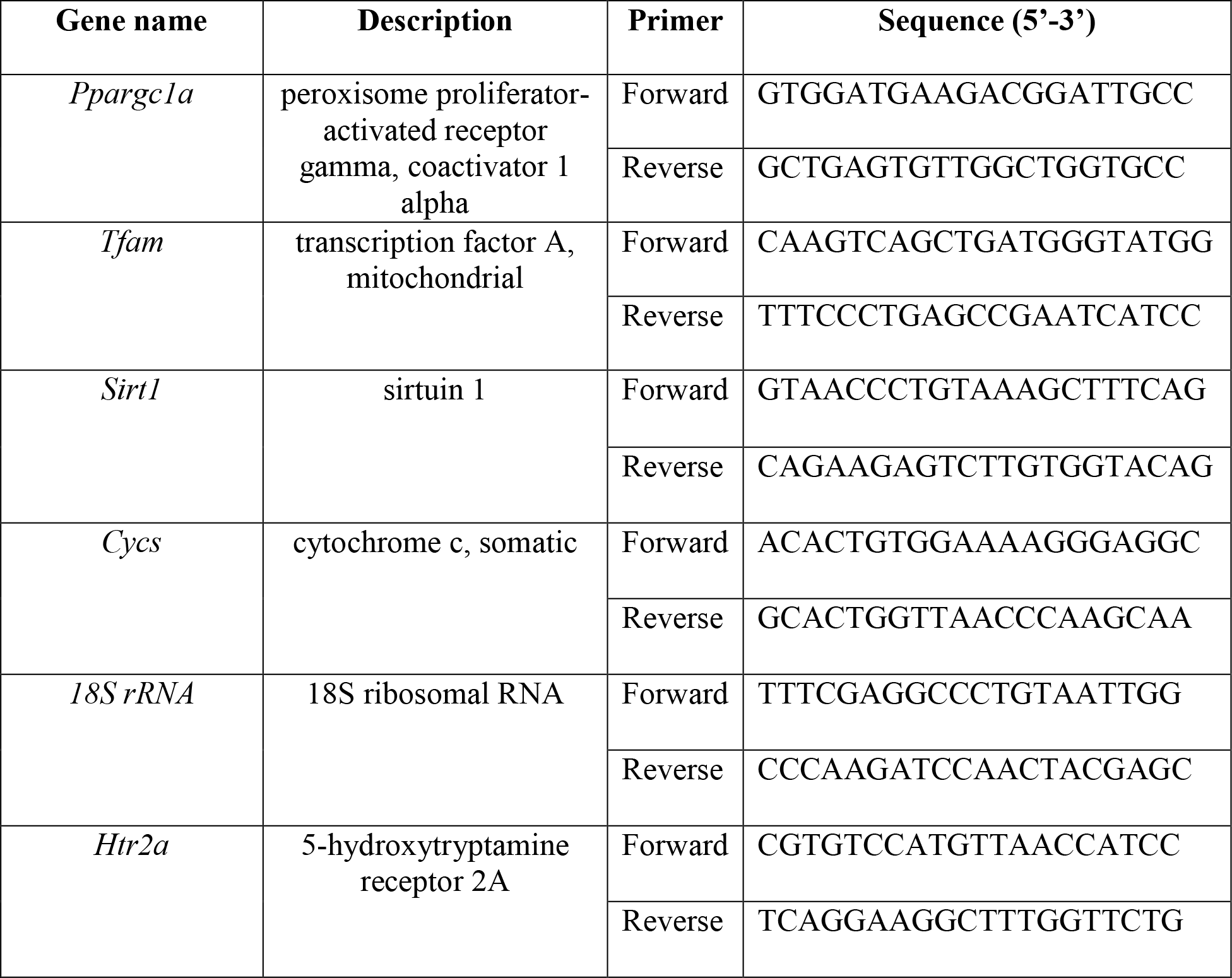
Primer sequences used for quantitative PCR analysis of mouse cDNA

**Table S4:**
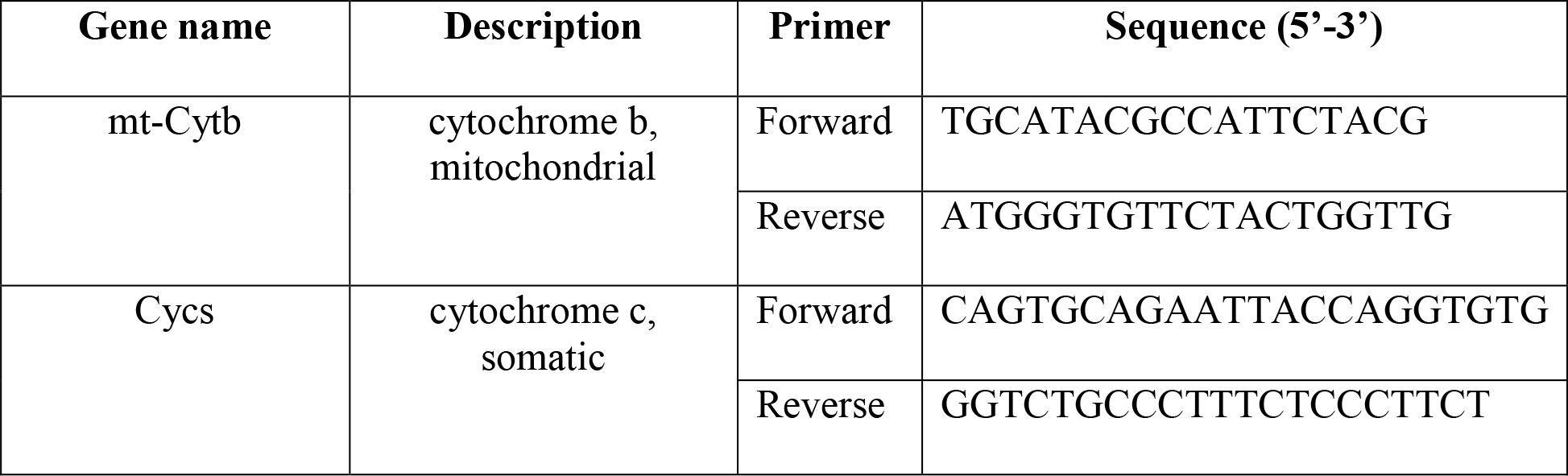
Primer sequences used for relative mitochondrial DNA (mtDNA) analysis in Mouse

## List of abbreviations

1. 1,4-Diamino-2,3-dicyano-1,4-bis[2-aminophenylthio]butadiene - U0126
2. 1-[6-[[(17β)-3-Methoxyestra-1,3,5(10)-trien-17-yl]amino]hexyl]-1H-pyrrole-2,5-dione - U73122
3. 2-(4-Morpholinyl)-8-phenyl-4H-1-benzopyran-4-one hydrochloride - LY294002
4. 4-iodo-2,5-dimethoxyphenylisopropylamine - DOI
5. 5-(and-6)-carboxy-2’,7’-dichlorodihydrofluorescein diacetate - carboxy-H_2_DCFDA
6. 6-Chloro-2,3,4,9-tetrahydro-1H-Carbazole-1-carboxamide - EX-527
7. Brain-derived neurotrophic factor - BDNF
8. Carbonylcyanide-p-trifluoromethoxyphenylhydrazone - FCCP
9. Dimethyl sulfoxide - DMSO
10. Extracellular-signal-regulated kinase - ERK (also called MAPK)
11. Hydrogen peroxide - H_2_O_2_
12. Microtubule Associated Protein 2 – MAP2
13. Mitochondrial DNA - mtDNA
14. Mitogen-activated protein kinase - MAPK or MAP kinase (also called ERK)
15. Mitogen-activated protein kinase kinase - MEK
16. N-[2-[4-(2-Methoxyphenyl)-1-piperazinyl]ethyl]-N-2-pyridinylcyclohexanecarboxamide - WAY100,635
17. Oxidative phosphorylation - OXPHOS
18. Phosphatidylinositol 3-kinase - PI3K
19. Phospholipase C - PLC
20. Protein kinase B (PKB) - Akt
21. R-(1)-a-(2,3-dimethoxyphenyl)21-[2-(4-flurophenylethyl)]24-piperidine-methanol - MDL100,907
22. Reactive oxygen species - ROS
23. Serotonin (5-hydroxytryptamine) - 5-HT

## References

1. Turlejski K (1996) Evolutionary ancient roles of serotonin: long-lasting regulation of activity and development. Acta NeurobiolExp (Wars) 56(2):619–36.

2. Berger M, Gray JA, Roth BL (2009) The Expanded Biology of Serotonin. Annu Rev Med 60(1):355–366.

3. Azmitia EC (2007) Serotonin and brain: evolution, neuroplasticity, and homeostasis. Int Rev Neurobiol 77:31–56.

4. Gaspar P, Cases O, Maroteaux L (2003) The developmental role of serotonin: News from mouse molecular genetics. Nat Rev Neurosci 4(12):1002–1012.

5. Lesch KP, Waider J (2012) Serotonin in the Modulation of Neural Plasticity and Networks: Implications for Neurodevelopmental Disorders. Neuron 76(1): 175–191.

6. Deneris E, Gaspar P (2018) Serotonin neuron development: shaping molecular and structural identities. Wiley Interdiscip Rev Dev Biol 7(1):11–16.

7. Dooley AE, Pappas IS, Parnavelas JG (1997) Serotonin Promotes the Survival of Cortical Glutamatergic Neurons in Vitro. Exp Neurol 148(1):205–214.

8. Mattson MP, Gleichmann M, Cheng A (2008) Mitochondria in Neuroplasticity and Neurological Disorders. Neuron 60(5):748–766.

9. Morava É, Kozicz T (2013) Mitochondria and the economy of stress (mal)adaptation. Neurosci Biobehav Rev 37(4):668–680.

10. Romanov RA, et al. (2018) Chemical synapses without synaptic vesicles: Purinergic neurotransmission through a CALHM1 channel-mitochondrial signaling complex. Sci Signal 11(529):eaao1815.

11. Mattson MP, Moehl K, Ghena N, Schmaedick M, Cheng A (2018) Intermittent metabolic switching, neuroplasticity and brain health. Nat Rev Neurosci 19(2):63–80.

12. Devine MJ, Kittler JT (2018) Mitochondria at the neuronal presynapse in health and disease. Nat Rev Neurosci 19(2):63–80.

13. Raefsky SM, Mattson MP (2017) Adaptive responses of neuronal mitochondria to bioenergetic challenges: Roles in neuroplasticity and disease resistance. Free Radic Biol Med 102(November 2016):203–216.

14. Hock MB, Kralli A (2009) Transcriptional Control of Mitochondrial Biogenesis and Function. Annu Rev Physiol 71(1):177–203.

15. Dominy JE, Puigserver P (2013) Mitochondrial Biogenesis through Activation of Nuclear Signaling Proteins. Cold Spring Harb Perspect Biol 5(7):a015008–a015008.

16. Wareski P, et al. (2009) PGC-1{alpha} and PGC-1{beta} regulate mitochondrial density in neurons. J Biol Chem 284(32):21379–85.

17. Scarpulla RC, Vega RB, Kelly DP (2012) Transcriptional integration of mitochondrial biogenesis. Trends Endocrinol Metab 23(9):459–466.

18. Rodgers JT, Lerin C, Gerhart-Hines Z, Puigserver P (2008) Metabolic adaptations through the PGC-1 alpha and SIRT1 pathways. FEBS Lett 582(1):46–53.

19. Finck BN, Kelly DP (2006) PGC-1 coactivators: inducible regulators of energy metabolism in health and disease. J Clin Invest 116(3):615–22.

20. Chen S, Owens GC, Crossin KL, Edelman DB (2007) Serotonin stimulates mitochondrial transport in hippocampal neurons. Mol Cell Neurosci 36(4):472–83.

21. Scholpa NE, Lynn MK, Corum D, Boger HA, Schnellmann RG (2018) 5-HT _1F_ receptor-mediated mitochondrial biogenesis for the treatment of Parkinson’s disease. Br J Pharmacol 175(2):348–358.

22. Brunet A, et al. (2004) Stress-dependent regulation of FOXO transcription factors by the SIRT1 deacetylase. Science 303(5666):2011–5.

23. St-Pierre J, et al. (2006) Suppression of reactive oxygen species and neurodegeneration by the PGC-1 transcriptional coactivators. Cell 127(2):397–408.

24. Rasbach K, Funk J, Jayavelu T, Green P, Schnellmann R (2010) {5-Hydroxytryptamine} Receptor Stimulation of Mitochondrial Biogenesis. J Pharmacol Exp Ther 332(2):632–639.

25. Harmon JL, et al. (2016) 5-HT2 Receptor Regulation of Mitochondrial Genes: Unexpected Pharmacological Effects of Agonists and Antagonists. J Pharmacol Exp Ther 357(1):1–9.

26. Garrett SM, Whitaker RM, Beeson CC, Schnellmann RG (2014) Agonism of the 5-Hydroxytryptamine 1F Receptor Promotes Mitochondrial Biogenesis and Recovery from Acute Kidney Injury. J Pharmacol Exp Ther 350(2):257–264.

27. Gibbs WS, et al. (2018) 5-HT1F receptor regulates mitochondrial homeostasis and its loss potentiates acute kidney injury and impairs renal recovery. Am J Physiol Physiol: ajprenal.00077.2018.

28. Nebigil CG, Etienne N, Messaddeq N, Maroteaux L (2003) Serotonin is a novel survival factor of cardiomyocytes:mitochondria as a target of 5-HT2B-receptor signaling. FASEB J 7(3): 1–24.

29. Braz GRF, et al. (2016) Neonatal SSRI exposure improves mitochondrial function and antioxidant defense in rat heart. Appl Physiol Nutr Metab 41(4):362–369.

30. Simões-Alves AC, et al. (2018) Neonatal treatment with fluoxetine improves mitochondrial respiration and reduces oxidative stress in liver of adult rats. J Cell Biochem 119(8):6555–6565.

31. Silva TLA, et al. (2018) Serotonin transporter inhibition during neonatal period induces sex-dependent effects on mitochondrial bioenergetics in the rat brainstem. Eur J Neurosci 48(1):1620–1634.

32. Van Der Kooij MA, et al. (2018) Diazepam actions in the VTA enhance social dominance and mitochondrial function in the nucleus accumbens by activation of dopamine D1 receptors. Mol Psychiatry 23(3):569–578.

33. Chen S, Owens GC, Edelman DB (2008) Dopamine inhibits mitochondrial motility in hippocampal neurons. PLoS One 3(7):e2804.

34. Zhou H, et al. (2014) N-acetyl-serotonin offers neuroprotection through inhibiting mitochondrial death pathways and autophagic activation in experimental models of ischemic injury. J Neurosci 34(8):2967–78.

35. Yang Y, et al. (2015) Melatonin prevents cell death and mitochondrial dysfunction via a SIRT1-dependent mechanism during ischemic-stroke in mice. J Pineal Res 58(1):61–70.

36. Xue F, et al. (2017) Melatonin Mediates Protective Effects against Kainic Acid-Induced Neuronal Death through Safeguarding ER Stress and Mitochondrial Disturbance. Front Mol Neurosci 10(February):1–16.

37. Daws LC, Toney GM (2007) High-Speed Chronoamperometry to Study Kinetics and Mechanisms for Serotonin Clearance In Vivo eds C. Borland A, M. Michael L (Boca Raton (FL): CRC Press/Taylor & Francis) doi:NBK2570[bookaccession].

38. Daubert EA, Condron BG (2010) Serotonin: a regulator of neuronal morphology and circuitry. Trends Neurosci 33(9):424–434.

39. Bunin MA, Wightman RM (1999) Paracrine neurotransmission in the CNS: involvement of 5-HT. Trends Neurosci 22(9):377–82.

40. Bunin MA, Wightman RM (1998) Quantitative evaluation of 5-hydroxytryptamine (serotonin) neuronal release and uptake: an investigation of extrasynaptic transmission. J Neurosci 18(13):4854–60.

41. Kaushalya SK, et al. (2008) Three-photon microscopy shows that somatic release can be a quantitatively significant component of serotonergic neurotransmission in the mammalian brain. J Neurosci Res 86(15):3469–3480.

42. Balaji J, Desai R, Kaushalya SK, Eaton MJ, Maiti S (2005) Quantitative measurement of serotonin synthesis and sequestration in individual live neuronal cells. J Neurochem 95(5):1217–26.

43. Riccio O, et al. (2009) Excess of serotonin affects embryonic interneuron migration through activation of the serotonin receptor 6. Mol Psychiatry 14(3):280–90.

44. Lavdas AA, Blue ME, Lincoln J, Parnavelas JG (1997) Serotonin promotes the differentiation of glutamate neurons in organotypic slice cultures of the developing cerebral cortex. J Neurosci 17(20):7872–80.

45. Kondoh M, Shiga T, Okado N (2004) Regulation of dendrite formation of Purkinje cells by serotonin through serotonin1A and serotonin2A receptors in culture. Neurosci Res 48(1):101–9.

46. Leopoldt D, et al. (1998) Gbetagamma stimulates phosphoinositide 3-kinase-gamma by direct interaction with two domains of the catalytic p110 subunit. J Biol Chem 273(12):7024–9.

47. Millan M, Marin P, Bockaert J, Mannourylacour C (2008) Signaling at G-protein-coupled serotonin receptors: recent advances and future research directions. Trends Pharmacol Sci 29(9):454–464.

48. González-Maeso J, et al. (2007) Hallucinogens Recruit Specific Cortical 5-HT2AReceptor-Mediated Signaling Pathways to Affect Behavior. Neuron 53(3):439–452.

49. Jones KA, et al. (2009) Rapid modulation of spine morphology by the 5-HT2A serotonin receptor through kalirin-7 signaling. Proc Natl Acad Sci U S A 106(46):19575–80.

50. Ly C, et al. (2018) Psychedelics Promote Structural and Functional Neural Plasticity. Cell Rep 23(11):3170–3182.

51. Vaidya VA, Marek GJ, Aghajanian GK, Duman RS (1997) 5-HT2A receptor-mediated regulation of brain-derived neurotrophic factor mRNA in the hippocampus and the neocortex. J Neurosci 17(8):2785–95.

52. Su B, Ji Y-S, Sun X, Liu X-H, Chen Z-Y (2014) Brain-derived neurotrophic factor (BDNF)-induced mitochondrial motility arrest and presynaptic docking contribute to BDNF-enhanced synaptic transmission. J Biol Chem 289(3):1213–26.

53. Cheng A, et al. (2012) Involvement of PGC-1a in the formation and maintenance of neuronal dendritic spines. Nat Commun 3(1):1250.

54. Price NL, et al. (2012) SIRT1 is required for AMPK activation and the beneficial effects of resveratrol on mitochondrial function. Cell Metab 15(5):675–90.

55. Philp A, et al. (2011) Sirtuin 1 (SIRT1) deacetylase activity is not required for mitochondrial biogenesis or peroxisome proliferator-activated receptor-gamma coactivator-1alpha (PGC-1alpha) deacetylation following endurance exercise. J Biol Chem 286(35):30561–70.

56. Rodgers JT, Puigserver P (2007) Fasting-dependent glucose and lipid metabolic response through hepatic sirtuin 1. Proc Natl Acad Sci U S A 104(31):12861–6.

57. Gerhart-Hines Z, et al. (2007) Metabolic control of muscle mitochondrial function and fatty acid oxidation through SIRT1/PGC-1alpha. EMBO J 26(7):1913–23.

58. Dorn GW, Vega RB, Kelly DP (2015) Mitochondrial biogenesis and dynamics in the developing and diseased heart. Genes Dev 29(19): 1981–1991.

59. Kelly DP, Scarpulla RC (2004) Transcriptional regulatory circuits controlling mitochondrial biogenesis and function. Genes Dev 18(4):357–68.

60. Libert S, et al. (2011) SIRT1 activates MAO-A in the brain to mediate anxiety and exploratory drive. Cell 147(7):1459–72.

61. Weger M, Sandi C (2018) High anxiety trait: A vulnerable phenotype for stress-induced depression. Neurosci BiobehavRev 87:27–37.

62. Picard M, McEwen BS, Epel ES, Sandi C (2018) An energetic view of stress: Focus on mitochondria. Front Neuroendocrinol 49(January):72–85.

63. Nicholls DG (2008) Oxidative Stress and Energy Crises in Neuronal Dysfunction. Ann N Y Acad Sci 1147(1):53–60.

64. Uittenbogaard M, Chiaramello A (2014) Mitochondrial Biogenesis: A Therapeutic Target for Neurodevelopmental Disorders and Neurodegenerative Diseases. Curr Pharm Des 20(35):5574–5593.

65. Cameron RB, Beeson CC, Schnellmann RG (2016) Development of Therapeutics That Induce Mitochondrial Biogenesis for the Treatment of Acute and Chronic Degenerative Diseases. J Med Chem 59(23):10411–10434.

66. Ben-Shachar D, Ene HM (2018) Mitochondrial Targeted Therapies: Where Do We Stand in Mental Disorders? Biol Psychiatry 83(9):770–779.

67. El-Hattab AW, Zarante AM, Almannai M, Scaglia F (2017) Therapies for mitochondrial diseases and current clinical trials. Mol Genet Metab 122(3):1–9.

68. Manji H, et al. (2012) Impaired mitochondrial function in psychiatric disorders. Nat Rev Neurosci 13(5):293–307.

69. Onyango IG, et al. (2010) Regulation of neuron mitochondrial biogenesis and relevance to brain health. Biochim Biophys Acta - Mol Basis Dis 1802(1):228–234.

70. Nierenberg AA, et al. (2018) Peroxisome Proliferator-Activated Receptor Gamma Coactivator-1 Alpha as a Novel Target for Bipolar Disorder and Other Neuropsychiatric Disorders. Biol Psychiatry 83(9):761–769.

71. Shao L, et al. (2008) Mitochondrial involvement in psychiatric disorders. Ann Med 40(4):281–295.

72. Li H, et al. (2007) SirT1 modulates the estrogen-insulin-like growth factor-1 signaling for postnatal development of mammary gland in mice. Breast Cancer Res 9(1):1–12.

## References

1. Banker G, Goslin K (1998) Culturing nerve cells. Cellular and Molecular Neuroscience Series, eds Banker G, Goslin K (MIT University Press,Cambridge, MA), pp 339–370. 2nd Ed.

2. Pacifici M, Peruzzi F (2012) Isolation and culture of rat embryonic neural cells: a quick protocol. J Vis Exp (63):e3965.

3. Kondoh M, Shiga T, Okado N (2004) Regulation of dendrite formation of Purkinje cells by serotonin through serotonin1A and serotonin2A receptors in culture. Neurosci Res 48(1):101–9.

4. Lavdas AA, Blue ME, Lincoln J, Parnavelas JG (1997) Serotonin promotes the differentiation of glutamate neurons in organotypic slice cultures of the developing cerebral cortex. J Neurosci 17(20):7872–80.

5. Riccio O, et al. (2009) Excess of serotonin affects embryonic interneuron migration through activation of the serotonin receptor 6. Mol Psychiatry 14(3):280–90.

6. Bunin MA, Wightman RM (1998) Quantitative evaluation of 5-hydroxytryptamine (serotonin) neuronal release and uptake: an investigation of extrasynaptic transmission. J Neurosci 18(13):4854–60.

7. Bunin MA, Wightman RM (1999) Paracrine neurotransmission in the CNS: involvement of 5-HT. Trends Neurosci 22(9):377–82.

8. Daws LC, Toney GM (2007) High-Speed Chronoamperometry to Study Kinetics and Mechanisms for Serotonin Clearance In Vivo eds C.Borland A, M.Michael L (Boca Raton (FL): CRC Press/Taylor & Francis) doi:NBK2570[bookaccession].

9. Schmittgen TD, Livak KJ (2008) Analyzing real-time PCR data by the comparative CT method. Nat Protoc 3(6):1101–1108.

10. Ghosh S, Banerjee KK, Vaidya VA, Kolthur-Seetharam U (2016) Early Stress History Alters Serum Insulin-Like Growth Factor-1 and Impairs Muscle Mitochondrial Function in Adult Male Rats. J Neuroendocrinol 28(9). doi:10.1111/jne.12397.

